# The life cycle of an archaeon with multiple membranes

**DOI:** 10.64898/2026.09.08.750209

**Authors:** Florian Mayer, Andriko von Kügelgen, Richard Stöckl, Veronika Seitz, Joe Parham, Zhexin Wang, Robert Reichelt, Tanmay A.M. Bharat, Harald Huber, Anja Spang, Dina Grohmann, Buzz Baum

## Abstract

Many prokaryotes are diderms. They divide using an FtsZ division ring to simultaneously constrict physically coupled inner and outer membranes to form daughter cells with two membranes. Currently, only one archaeon, *Ignicoccus hospitalis*, is known to have an outer and inner membrane. Here, in exploring *Ignicoccus* cell division, we show by expansion microscopy that *I. hospitalis* uses a contractile CdvA-ESCRT-III ring to repeatedly constrict and cut its inner membrane in a way that yields clusters containing multiple cytoplasms within a shared periplasm. The outer bounding membrane then bursts to release the progeny cells, which regain their parental cell architecture via a membrane duplication process. Furthermore, this life cycle appears conserved across *Ignicoccus* strains, even though the precise timing of events changes. Taken together, these data show that *Ignicoccus* cells propagate by undergoing dynamic changes in their architecture as they transit between single and multi-cellular states.

## Introduction

Some bacteria possess a single bounding plasma membrane whereas others including *Escherichia coli* are bounded by two membranes (*1*). Surprisingly, it has recently become clear that the ‘diderm’ state is ancestral (*2*). In most of these bacteria, the outer membrane fulfils a structural role (*3*) acting as a physical barrier that protects cells from the environment, whereas the inner membrane separates the chemistry of the cytoplasm from the external world, and sustains ionic gradients. In a typical diderm, like *E. coli*, the two membranes are mechanically tethered by physical links between them, including those mediated through the peptidoglycan cell wall (*4*). As a result, when the cells divide, via a process that is typically guided by a cytoplasmic FtsZ ring (*5*) and cell wall assembly (*6*), the two membranes are remodelled together. This ensures that the diderm membrane architecture of the parental cell is preserved through division to yield daughter cells, each of which has two bounding membranes. This completes the cell’s life cycle.

In contrast to bacteria, nearly all archaea, including recently cultivated members of the Asgard archaea (*7–9*), officially referred to as *Asgardarchaeota* (*10*) and/or *Prometheoarchaeota* (*11*) are bounded by a single membrane and, with few exceptions (*12*), lack a cell wall (*13*). Because of this, most protect themselves from the environment via a highly ordered surface lattice of closely packed glycosylated proteins - often termed the S-layer (*13*, *14*).

While the genomes of most archaea (including the Asgard archaea (*15–20*)) encode homologues of FtsZ (*21*), which drive division in the vast majority of bacteria (*22*), FtsZ is lacking in the genomes of many members of the archaeal phylum *Thermoproteota*, such as *Sulfolobus* (*21*). In these strains, cells tend to be spherical, and divide using a contractile ring that relies on the protein CdvA acting together with proteins related to eukaryotic SNF7 domain proteins as part of ESCRT-III (**E**ndosomal **S**orting **C**omplex **R**equired for **T**ransport) and Vps4 (*23–25*). Strikingly, ESCRT-III proteins are homologous to the machinery that has been shown to constrict and cut membranes at the end of cytokinesis in human cells (*26*).

Cytokinesis in the monoderm *Sulfolobus,* where this mode of division is probably best understood within archaea (*23–25*), begins with the formation of a CdvA ring at the centre of the spherical cell (*27*). CdvA then recruits a set of ESCRT-III homologues (referred to as CdvB proteins), which form a division ring whose composition changes as it constricts as the result of Vps4 (CdvC)-dependent polymer disassembly (*23*).

Like the *Sulfolobales*, the genome of *Ignicoccus hospitalis* lacks FtsZ and actin homologues. Instead, its genome encodes a single CdvA protein (gene Igni_0996), three ESCRT-III homologues, namely CdvB (gene Igni_0995), CdvB1 (gene Igni_1156), CdvB2 (gene Igni_0101), and one Vps4 (CdvC) protein (gene Igni_0994). Phylogenetics, protein structure prediction and structural alignments of CdvB, CdvB1 and CdvB2 (Figures S1 and S2) suggest that *Sulfolobus* and *Ignicoccus* have a similar division machinery. However, despite having genomes that encode a similar division machinery, *Ignicoccus* cells differ from *Sulfolobus* and all other archaeal species described so far in having a unique membrane architecture. *Ignicoccus* cells possess a convoluted inner membrane, and a smooth bounding outer membrane (Figure 1A) (*28–31*). Because *Ignicoccus* cells also lack an S-layer, it is likely that the outer membrane provides protection from the environment (*29*). Note that *I. hospitalis* is a hyperthermophilic archaeon with an optimal growth themperature of 90°C (*28*). This raises the question as to how *Ignicoccus* cells are able to use their conserved CdvA-ESCRT-III division machinery to divide in a manner that preserves their complex cellular architecture, with two membranes, across multiple generations.

**Figure 1.**
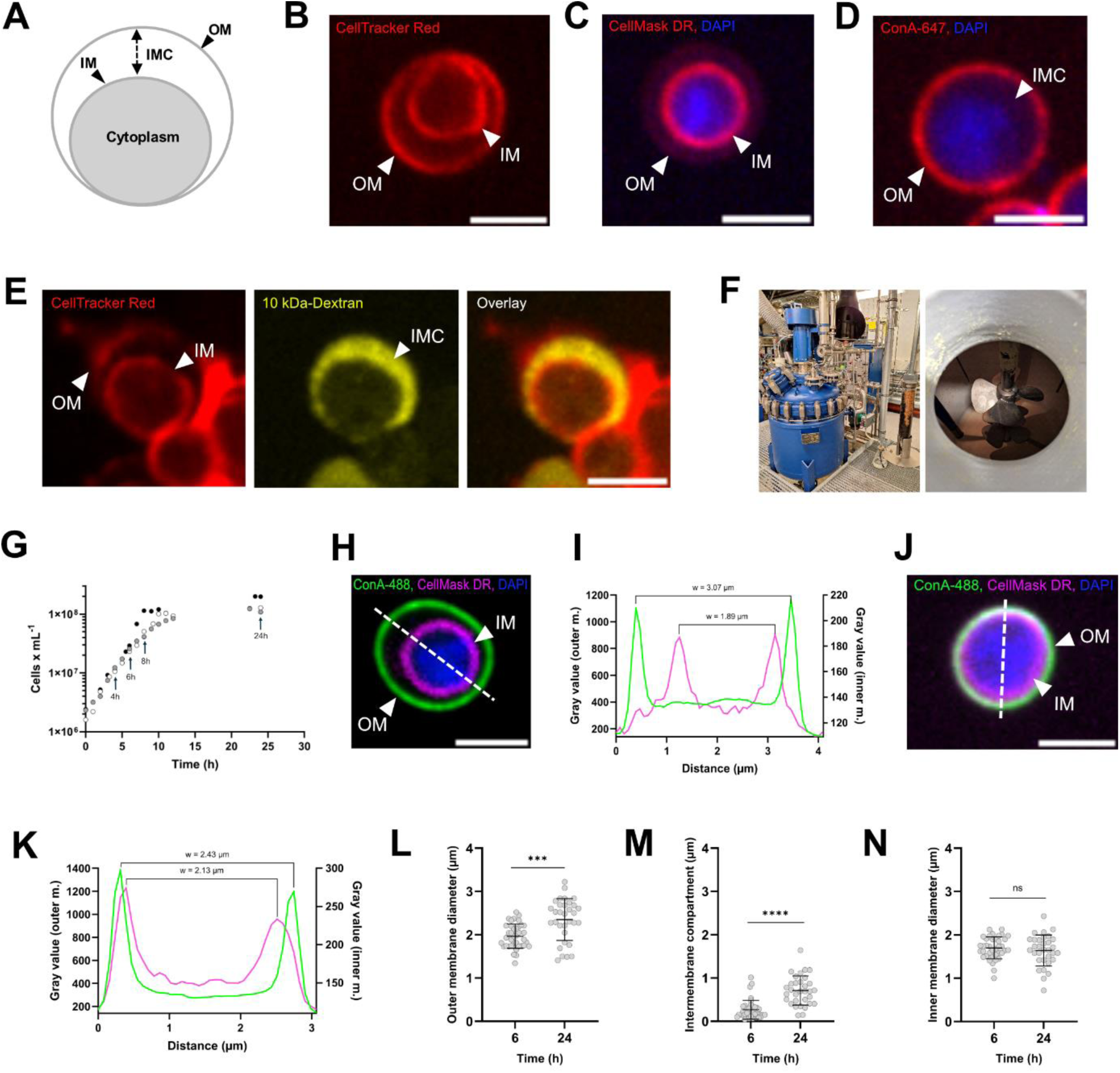
Cell architecture of *Ignicoccus hospitalis* during growth. **A,** Schematic showing the cell architecture of *I. hospitalis* with outer membrane (OM), inner membrane (IM), the inter-membrane compartment (IMC) and cytoplasm. **B,** Staining of both membranes with CellTracker Red CMTPX. Scale bar, 2 µm. **C,** Staining of the inner membrane with CellMask Deep Red and the DNA in the cytoplasm with DAPI. Scale bar, 2 µm. **D,** Staining of the outer membrane with the lectin ConA-647 and the DNA in the cytoplasm with DAPI. Scale bar, 2 µm. **E,** Staining of the inter-membrane compartment with 10 kDa Dextran Oregan Green, labelling of membranes with CellTracker Red CMTPX. Scale bar, 2 µm. **F,** 300 L bioreactor with steering unit used to cultivate *I. hospitalis* at 90°C and with a continuous H2/CO2 gas flow. **G,** Growth curve of *I. hospitalis* cultivated in the 300 L bioreactor in three replicates (black, grey and white dots). Cells for analyses were taken at 4, 6, 8 and 24 h after inoculation. **H,** Multicolor cell labelling with ConA-488 staining the outer membrane, CellMask Deep Red staining the inner membrane and DAPI to stain DNA. Scale bar, 2 µm. Dotted line represents the diagonal used for cell measurements of a stationary phase cell shown in panel I. **I,** Measurement of the diameter of outer and inner membrane from a stationary phase cell shown in panel H. The diameters were determined by measuring the distance (width) between peak-to-peak of a line profile. Diameter of outer membrane (green) and diameter of inner membrane (magenta). **J,** Mid-log phase cell, staining of outer membrane with ConA-488 and inner membrane with CellMask Deep Red. DNA was stained with DAPI. Scale bar, 2 µm. Dotted line represents the diagonal used for cell measurements of a mid-log phase cell shown in panel K. **K,** Measurement of the diameter of inner and outer membrane of mid-log phase cell shown in panel J. The diameters were determined by measuring the distance (width) between peak-to-peak of a line profile. Diameter of outer membrane (green) and diameter of inner membrane (magenta). **L,** Significant change of the outer membrane diameter between cells from mid-log- and stationary phase (6 h vs. 24 h). *** = *p*-value = 0.0001, Mann-Whitney U test. **M,** Significant change of the intermembrane compartment size between cells from mid-log- and stationary phase (6 h vs. 24 h). **** = *p*-value < 0.0001, Mann-Whitney U test. **N,** No significant change of inner membrane diameter for cells at mid-log- and stationary phase (6 h vs. 24 h). ns = *p*-value = 0.5130, Mann-Whitney U test.

To address this question, we employ fluorescent cell imaging without fixation, expansion microscopy and cryogenic electron microscopy. Our analysis revealed that *I. hospitalis* first forms a CdvA and ESCRT-III based division ring inside the inner cell membrane. Subsequently, the ring constricts and cuts the inner cell membrane to generate a cell cluster that contains two cytoplasms within a single periplasmic space bounded by an outer membrane. As this process is repeated, clusters are formed that burst to release multiple progeny cells, whose double membrane architecture is restored via membrane duplication, enabling them to complete this lifecycle. Importantly, we show that this unusual life cycle is broadly conserved across the *Ignicoccus* genus, though lineage-specific differences in the timing of membrane duplication were observed. Taken together, this work identifies a novel mode of cell propagation.

## Results

Members of the genus *Ignicoccus* are the only well-studied archaea known to possess two membranes separated by a wide intermembrane compartment (Figure 1A) (*28*). Given this unusual cell architecture, and the lack of a cell wall or S-layer, it is unclear how *Ignicoccus* cells divide in a manner that preserves their architecture across generations. Studying the mechanism of *I. hospitalis* cell division is extremely challenging because these cells are small, with a size of 1-4 µm, and grow at 90 °C under strictly anaerobic conditions using hydrogen, carbon dioxide and elemental sulphur to support their chemolithoautotrophic metabolism.

To study cell division in this system, our first task was to develop tools to investigate their cell architecture. To do so, we harvested cells from growing cultures and rapidly cooled them to ambient temperatures without fixation - forcing cells into a state of suspended animation without loss of viability as measured by efficient growth after reinoculation into fresh medium (Figure S3). By screening for membrane dyes (*32*) that stain the inner and/or outer membrane of unfixed *I. hospitalis* cells, we identified CellTracker Red CMTPX as a label that marks both the outer and inner membrane (Figure 1B), while CellMask Deep Red preferentially labels the inner cell membrane (Figure 1C). We were also able to use DAPI as a DNA stain (Figure 1C), and a fluorescently labeled lectin Concanavalin A (ConA) to label glycosylated lipids (*33*) that decorate the outer surface of non-permeabilized unfixed *I. hospitalis* cells (Figure 1D). In parallel, we were able to selectively label the intermembrane compartment (IMC) by adding fluorescently-labeled 10 kDa Dextran to cells at room temperature (Figure 1E), under conditions in which 70 kDa Dextran was excluded from cells entirely (Figure S4). This result implies that the outer membrane of *Ignicoccus* is permeable to certain complex hydrophilic molecules, similar to the outer membrane of Gram-negative bacteria (*34*).

### *I. hospitalis* cell architecture changes throughout growth phases

We were then able to use these selective membrane staining tools to study the architecture of unfixed cells taken from a 300 L bioreactor (Figure 1F) in which cells were grown under continuous flow of H2/CO2 gas and with sulphur as electron acceptor. Note that cultivation in a bioreactor was critical to obtain sufficient cell material at different time points across the growth cycle for the downstream analysis. After inoculation, cell numbers in the bioreactor increased in a reproducible manner (N=3), as measured by cell counts using brightfield microscopy (Figure 1G), before reaching stationary phase after about 24 hours. For the cell biological analysis, we sampled cells 4 hours after inoculation in early log phase, at 6 and 8 hours in mid-log phase, and at 24 hours, which represents stationary phase and marks the end of the cultivation (Figure 1G).

By imaging *I. hospitalis* cells at room temperature, we were able to measure cell size, inner membrane diameter and the maximum width of the inter-membrane compartment using the dyes ConA and CellMask Deep Red for differential membrane staining (Figures 1H and 1I), while cells remained in a state of suspended animation (Figure S3). Interestingly, when imaged in this way, most cells in the exponential phase of growth had a thin IMC or appeared to lack one altogether (Figures 1J and 1K). Even when two membranes could be discerned, the size of the IMC was often limited in exponentially growing cells. This changed as single cells reached stationary phase. In cells harvested at stationary phase, the size of the cells increased markedly (Figure 1L) due to a significant increase of the intermembrane compartment (Figure 1M), enabling the two membranes to be readily distinguished. The size of the cytoplasmic compartment, as measured by the inner membrane diameter, however, did not change (Figure 1N). These data suggest that the characteristic *I. hospitalis* architecture previously described in the literature (*28*, *30*, *31*, *35*, *36*), in which cells possess two well separated membranes and a large intermembrane compartment, predominantly reflects the architecture of cells in stationary phase.

### Immunolabeling and expansion microscopy reveal a new mode of cell division in *I. hospitalis*

In previous work, the *I. hospitalis* genome was reported to possess a CdvA-ESCRT-III division machinery, like that observed in *Sulfolobales* (*25*, *37*). To determine the growth phase-dependent expression of this machinery, we performed quantitative proteomics by mass spectrometry on cells harvested from bioreactor cultures during logarithmic and stationary growth phases. We used linear modelling to test for changes in protein abundances greater than 1.5-fold between growth phases, identifying 192 differentially abundant proteins at a 5% false discovery rate (Figure 2A). The previously characterized cell division proteins CdvA, CdvB and CdvB1 were most abundant during logarithmic phase and were significantly less abundant in stationary phase (Figure 2B). To validate the proteomics data set, we considered control proteins such like Ihomp1, an abundant outer membrane pore-forming protein (*38*). Ihomp1 showed no significant change in abundance across the growth cycle (Figure 2B). Another control, the type-4 pilin protein (gene Igni_0670) was significantly more abundant in stationary phase (Figure 2B).

**Figure 2.**
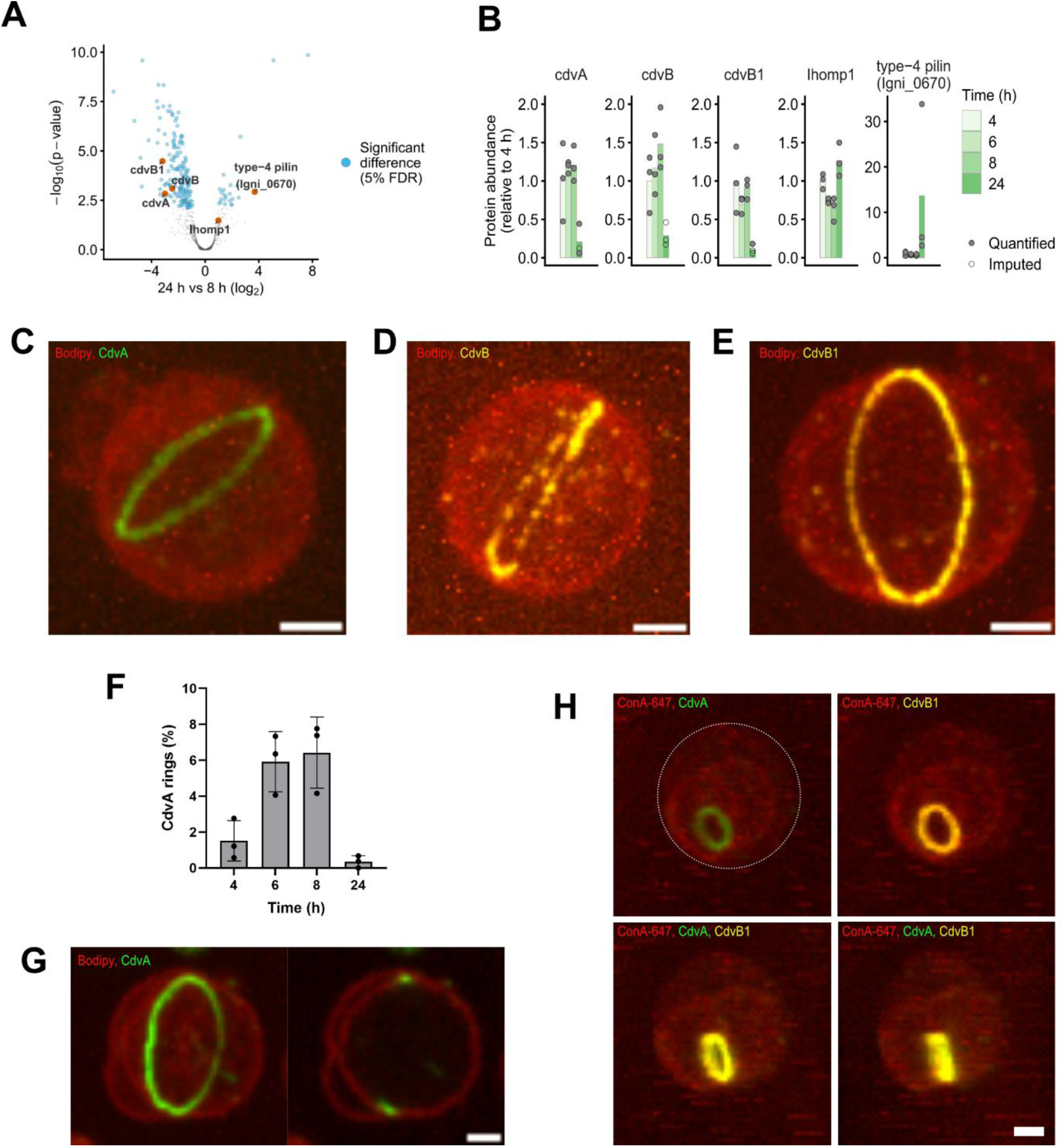
Cell division in *I. hospitalis* by CdvA and the ESCRT-III proteins CdvB and CdvB1. **A,** Linear modelling was used to test for changes in protein abundances between log (8 h) and stationary phase (24 h) cells of *I. hospitalis*, identifying 192 differentially abundant proteins at a 5% false discovery rate. **B,** Abundance of CdvA, CdvB, CdvB1, Ihomp1 and type-4 pilin (Igni_0670) proteins across the growth phases. Protein abundance is expressed relative to 4 h. Open and filled points denote imputed and directly quantified values, respectively. **C,** ExM was used to immunolocalize the CdvA ring (green) in *I. hospitalis* stained with Bodipy TR Ceramide (red). Maximum intensity projection. Scale bar, 2 µm. **D,** CdvB ring (yellow) of *I. hospitalis* (red) shown by ExM. Membranes were stained with Bodipy TR Ceramide. Maximum intensity projection. Scale bar, 2 µm. **E,** Immunolocalization of the CdvB1 ring (yellow) by ExM in *I. hospitalis* (red, stained with Bodipy TR Ceramide), shown in maximum intensity projection. Scale bar, 2 µm. **F,** Distribution of CdvA rings during the cell cycle of *I. hospitalis*. Highest abundance of CdvA rings at mid-log phase of growth. N=3, bioreactor cultivations. **G,** CdvA binds only to the inner membrane of *I. hospitalis* and forms a polymeric ring, as shown by ExM. Left picture, maximum intensity projection. Right picture one slide through the cell. CdvA (green) and membrane staining (red) with Bodipy TR Ceramide. Scale bar, 2 µm. **H,** CdvA and CdvB1-rings are involved in cell division and localized at the constriction site of two dividing *I. hospitalis* cells, as shown by ExM. White dotted line marks the outer membrane of the parental cell. CdvA ring (green), CdvB1 ring (yellow), cell stained with ConA-647 (red). Different views of cell cluster shown in maximum intensity projection. Scale bar, 2 µm.

To visualise division rings in *I. hospitalis* cells, we generated antibodies against CdvA, CdvB, and CdvB1 homologs. Due to the small size of *Ignicoccus* cells, the impermeability of archaeal membranes for high molecular weight molecules like antibodies, and the fact that membranes of *Ignicoccus* are disrupted by detergents even when fixed, a new approach was needed. We developed an expansion microscopy (ExM) protocol for immunofluorescence imaging (Figure S5). In doing so, embedding of cells into a gel and crosslinking of proteins preserved the structural integrity of the cell and cell expansion enabled permeabilization for antibody penetration (Figures 2C). At the same time, it also enabled us to physically enlarge cells (Figures S5H and S5I), increasing the resolution at which we could image division components. Using antibody-labeling and ExM in this way, we were able to visualise cell division rings in *I. hospitalis* cells labeled for CdvA (Figure 2C), CdvB (Figure 2D) and CdvB1 (Figure 2E); while also visualising the inner and outer membranes using Bodipy TR Ceramide or ConA. Since the CdvA antibody gave the most robust signal in cells at different stages of the division process (*39*), this marker was used to quantify division rings. By counting the number of CdvA rings in the population, we observed that division peaked during mid log-phase (Figure 2F). This finding is congruent with mass spectrometry measurements in which total CdvA levels reached their maximum during log-phase (Figure 2B).

Next, to ascertain how *Ignicoccus* cells with two membranes divide, we examined CdvA rings in cells in which both an inner and outer membrane could be clearly discerned using ExM. In every case, the CdvA ring was localized on the inside of the inner cell membrane (Figure 2G). These data support the idea that the division rings, which are an average of 9% (*n*=7) smaller than the outer membrane of the cell, bind and deform the inner membrane. Moreover, in the small number of cells in which CdvA and CdvB1 rings could be imaged mid-constriction, division rings pulled the inner membrane away from the spherical outer cell membrane (Figure 2H).

If, as suggested by these images, it is only the inner membrane of these cells that undergoes Cdv-dependent cytokinesis, one would expect *I. hospitalis* cells, that have completed division, to possess multiple cytoplasms within a single bounding membrane. This was indeed the case. Cell clusters containing multiple cytoplasms were observed both by light microscopy (Figures 3A and 3B) and cryogenic focused ion beam milling and scanning electron microscopy (cryo-FIB-SEM) (Figure 3C). When cells were stained with CellMask Deep Red to mark the inner membranes, fluorescently labeled ConA to highlight the outer membrane of cells, and DAPI to label the DNA, we observed multiple instances of clusters containing two or more cytoplasms contained within a single spherical outer membrane (Figures 3A and 3B). In addition, using ExM to increase the resolution of imaging we were able to observe clusters in which one of the two cytoplasms possessed a CdvA division ring that was attached to the inner membrane, but which had completely separated from the outer membrane (Figure 3D). Taken together, these data indicate that in growing cultures of *I. hospitalis*, the cytokinetic machinery selectively divides the inner cell membrane to generate clusters with multiple internal cells contained within a single bounding membrane.

**Figure 3.**
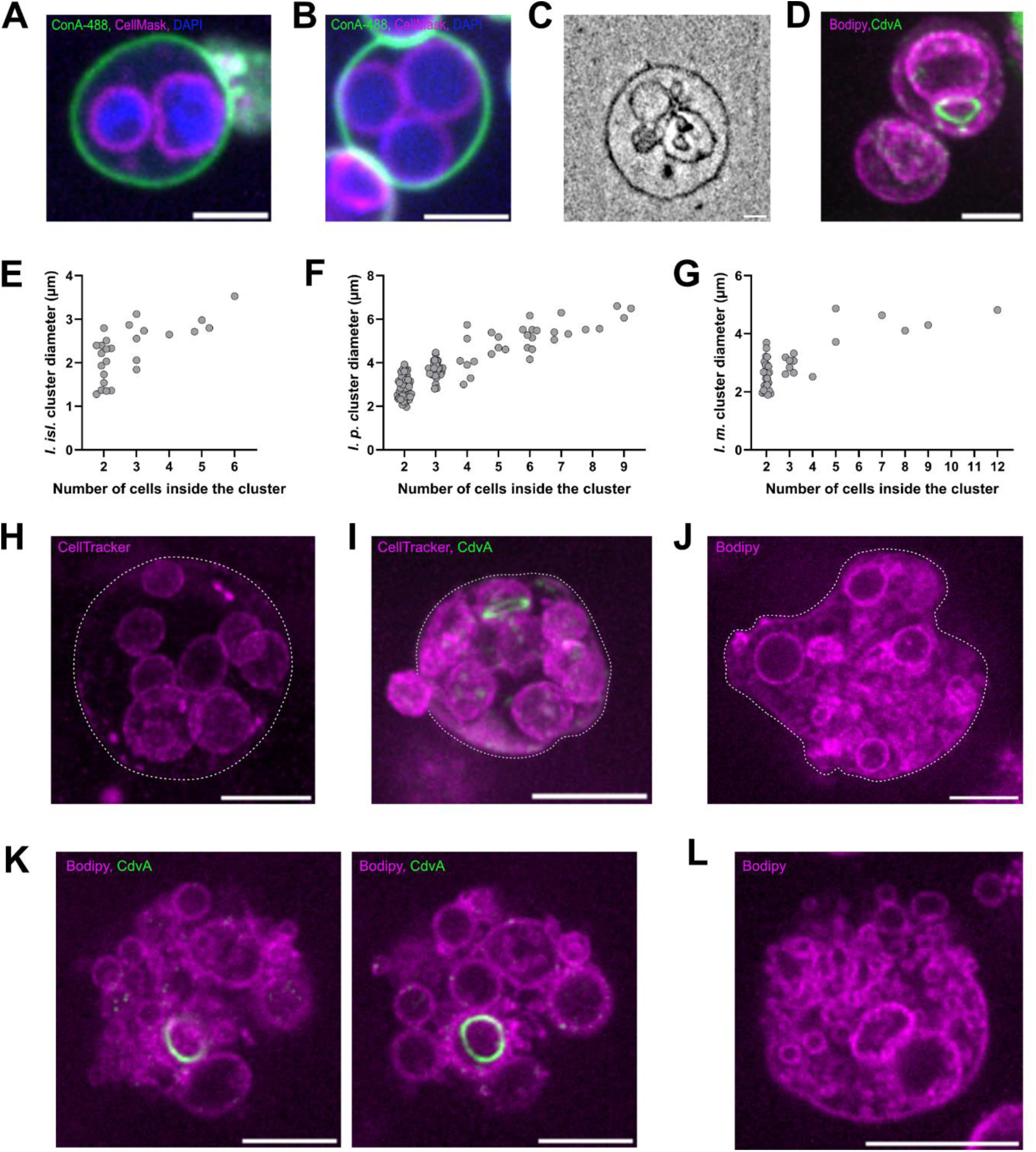
Cluster formation during cell division in different *Ignicoccus* strains. **A,** *I. hospitalis* cell stained with ConA-488, CellMask Deep Red and DAPI, with two cytoplasms bounded by a non-constricted outer membrane. Cells were visualized in agarose pads. Scale bar, 2 µm. **B,** Cell division in *I. hospitalis* leads to cell cluster formation. Clusters are bounded by a single outer membrane. Outer membrane stained with ConA-488, inner membrane with CellMask Deep Red and DNA with DAPI. Cells were visualized in a Fluoro dish. Scale bar, 2 µm. **C,** Cryo-FIB-SEM of *I. hospitalis* showing several inner cytoplasms bounded by an outer membrane. Scale bar, 250 nm. **D,** Immunolocalization of CdvA in *I. hospitalis* cell cluster by ExM. Cells stained with Bodipy TR Ceramide. Z-projection, scale bar, 5 µm. **E,** Diameter of *I. islandicus* cell cluster stained with CellTracker Red (both membranes) and MitoTracker Green (only inner membrane) and visualized on poly-L-lysine coated Fluoro dishes. **F,** Diameter of *I. pacificus* cell cluster stained with CellTracker Red and MitoTracker Green and visualized on agarose pads. **G,** Diameter of cell cluster of *Ca.* ‘Ignicoccus morulus’ stained with CellTracker Red and MitoTracker Green and visualized on ConA-coated Fluoro dishes. **H,** ExM of *I. pacificus* cell cluster stained with CellTracker Red, Z-projection, scale bar, 10 µm. **I,** *I. pacificus* cell cluster with CdvA ring visualized by ExM. Membranes stained with CellTracker Red and CdvA-488 staining with the CdvA antibody from *I. hospitalis*. Z-projection, scale bar, 10 µm. **J,** ExM of cell cluster of *I. hospitalis* with outer parental membrane from mid-log-phase of growth stained with Bodipy TR Ceramide. Scale bar, 10 µm. **K,** ExM of cell cluster of *I. hospitalis* showing no parental outer membrane. Cells from mid-log-phase of growth stained with Bodipy TR Ceramide and immunolocalization of CdvA. Two slides of a stack are shown. Scale bar, 10 µm. **L,** Cell cluster of *I. hospitalis* with disrupted outer membrane on one side and release of progeny cells. Scale bar, 10 µm.

### Conserved cell division mode in the genus *Ignicoccus*

To determine whether this unusual mode of cell division is conserved across the genus *Ignicoccus*, we imaged unfixed cells at room temperature from other *Ignicoccus* species using CellTracker Red to stain both membranes and MitoTracker Green FM to stain the inner membrane. Due to different membrane properties of the strains, the imaging approaches were slightly adapted to enable imaging under the best individual mounting conditions in each case (see methods). In *Ignicoccus islandicus*, which we imaged on poly-L-lysine-coated Fluoro dishes, we saw cell clusters containing 2 to 6 physically separate inner cytoplasmic compartments (Figure 3E). *Ignicoccus pacificus* clusters were imaged on agarose pads and contained between 2 and 9 compartments within a single bounding membrane (Figure 3F). Finally, we also examined a new isolate, *Candidatus ‘*Ignicoccus morulus*’*, which, as revealed by phylogenetic analysis, belongs to the *Ignicoccus* genus (Figure S6). When this new isolate was imaged on ConA-coated Fluoro dishes, clusters were observed containing between 2 and 12 internal compartments (Figure 3G).

Since *Ignicoccus* clusters in these cultures often contained an odd number of cytoplasmatic compartments (Figures 3E, 3F and 3G), it seems likely that these are generated by non-synchronous divisions. To test this, we employed the ExM procedure and the CdvA antibody of *I. hospitalis* to immunolocalize CdvA in *I. pacificus.* This strain had the highest number of cell clusters of all tested *Ignicoccus* strains (Figures 3E, 3F and 3G), could be easily expanded (Figure 3H) and could be labeled using the CdvA antibody we raised against the *I. hospitalis* protein (Figure 3I). In every case, we only observed a single individual cytoplasmic compartment undergoing cytokinesis within a large cluster (Figure 3I). This shows that cell division in *Ignicoccus* is asynchronous.

### Completion of cytokinesis through outer membrane rupture

When we quantified the number of internal compartments per cluster in *I. islandicus*, *I. pacificus* and *Ca.* ‘I. morulus’, we noted that the maximum size of clusters reached a plateau as the number of inner cytoplasms increased (Figures 3E, 3F and 3G). This raised the question as to whether the large clusters in *Ignicoccus* burst once the parental outer membrane of the cluster reached its limit. These clusters included some with multiple compartments inside a single bounding parental outer membrane (Figure 3J) as well as clusters that lacked a large bounding parental (Figure 3K, note the CdvA ring in one cell in the cluster that has a single bounding membrane).

Strikingly, we also observed clusters that appeared to be midway through the process of rupturing to release inner cells (Figure 3L). Taken together, these data imply that the *Ignicoccus* life cycle is completed and restarts anew when the outer membrane bursts.

### Formation of a new outer membrane

Nevertheless, an unresolved question remains. Completing the life cycle and restoring parental membrane organisation also requires the regeneration of a second membrane in cells emerging from a daughter cell with a single bounding membrane. To better understand how this is achieved, we focused our analysis on single isolated *Ignicoccus* cells which are not present in a cluster. We analyzed cells of *I. hospitalis* by electron cryotomography (cryo-ET) and cryo-FIB-SEM, showing that some cells of *I. hospitalis* indeed have only a single membrane bounding the ribosome-rich cytoplasm (Figure S7A and S7B). Moreover, cells with two membranes and an intermembrane compartment were found (Figure S7C and S7D). The same cell architecture was also observed with live cell imaging of non-fixed *I. hospitalis* cells (Figures 1B, 1E and 1H). In addition, as shown by cryo-FIB-SEM, also *I. pacificus* cultures contained cells with a single bounding membrane (Figure S7E) as well as cells with two membranes (Figure S7F).

Strikingly, we observed examples of *I. hospitalis* cells bounded by one membrane that have begun to split into two at two three-way junctions to generate a bubble-like periplasm in between (Figure 2G). To understand how membrane proteins might be partitioned and distinct membrane identity is generated as cells undergo the process of double membrane formation, we used antibodies to examine the localisation of AMP-forming Acetyl-CoA synthetase (ACS), a protein previously shown to be localised to the outer membrane (*40*). Interestingly, ACS antibodies localised to the membrane of isolated *I. hospitalis* cells that have a single bounding membrane (Figure 4A). In a subset of cells mid-way through the process of membrane duplication, which contain an intermembrane compartment blister, ACS localized to the outer membrane in regions where the two membranes could be readily distinguished (Figures 4B and 4C). Similarly, ACS antibodies nearly exclusively labeled the outer membrane of *I. hospitalis* cells and clusters that contain inner and outer membranes (Figures 4D and 4E). These data suggest that cells are able to sort proteins destined to be localised in the outer membrane during the process of membrane duplication, either via clearance of the protein from the inner membrane and/or via its re-distribution to the outer membrane.

**Figure 4.**
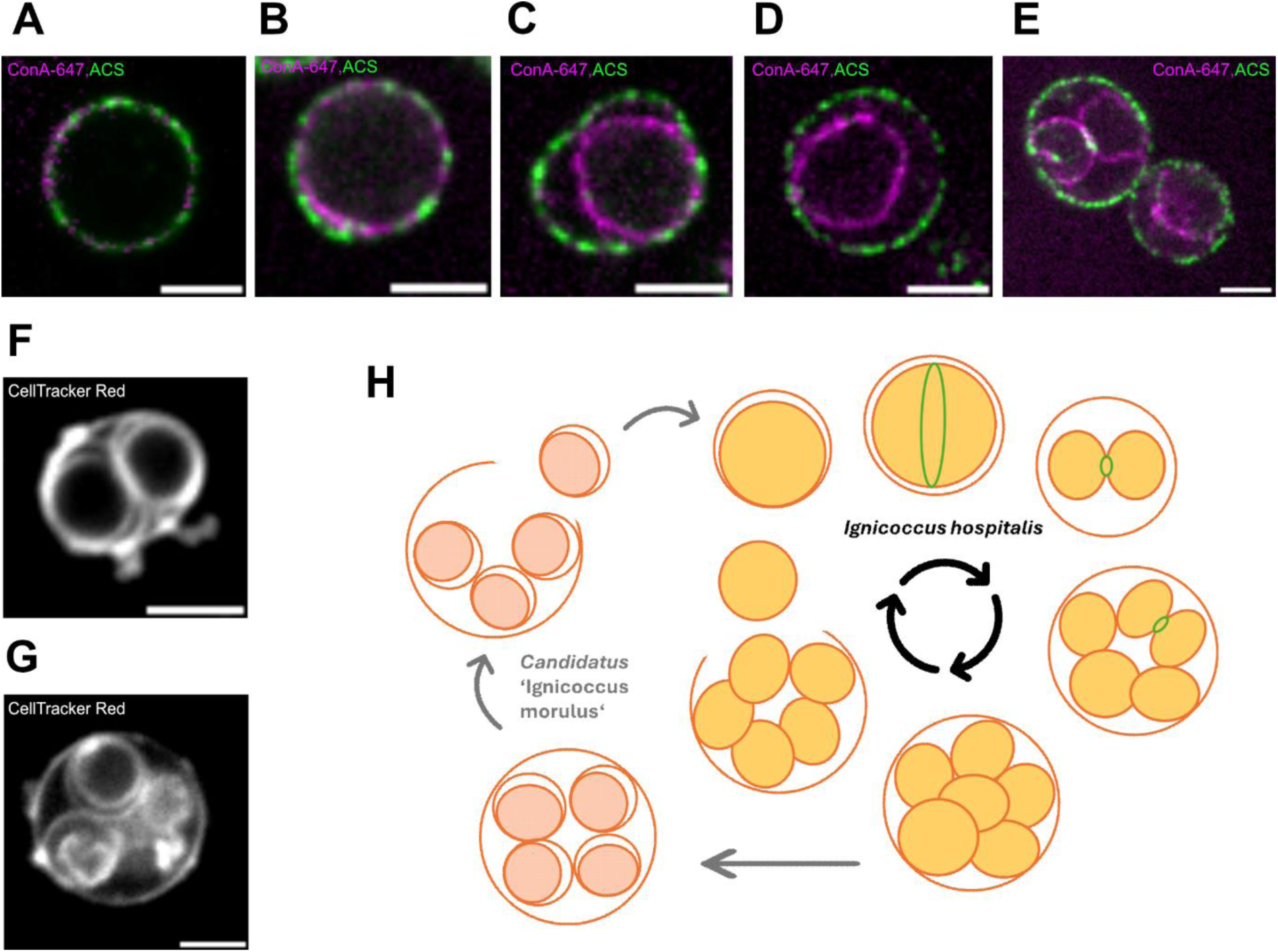
Membrane duplication in *Ignicoccus* progeny. **A to F,** ExM of *I. hospitalis* cells stained with ConA-647 and the acetyl-CoA synthetase (ACS) antibody coupled to AlexaFluor 488. Scale bars, 5 µm. ACS is a marker for the outer membrane. **A,** Cell with only one membrane show ACS localization in this membrane. **B,** Cells in the process of membrane duplication. ACS is localized in the outer membrane where the membrane starts to duplicate. **C,** The membrane duplicates at two independent blisters. Between both blisters the outer membrane is still connected to the inner membrane. ACS is mainly found in the outer membrane at this stage. **D,** Cells showing a clear outer membrane, IMC and inner membrane. At this stage, ACS is localized in the outer membrane only. **E,** During cell division, ACS is localized in the non-divided outer membrane of the cell cluster. Less ACS labelling is in the membranes of dividing daughter cells within the cluster. **F,** *Ca.* ‘I. morulus’ stained with CellTracker Red and visualized on ConA-coated Fluoro dishes. For the two divided cells inside the cell cluster, a second membrane can be seen leading to a cell cluster with three membranes in total. Scale bar, 2 µm. **G,** One slide of an image stack with *Ca.* ‘I. morulus’ stained with CellTracker Red and visualized on ConA-coated Fluoro dishes. The cell cluster contains five cells bounded by a single outer membrane. Three of the fives cells inside already formed a second membrane. Scale bar, 2 µm. **H,** Life cycle of *I. hospitalis* and *Ca.* ‘I. morulus’. Cell division rings (green) divide only the inner membrane, yielding two cytoplasm with a single bounding outer membrane. This process is repeated several times without division of the outer membrane leading to the formation of a cell cluster bounded by the parental outer membrane. When the outer membrane reaches its growth limit, it ruptures and releases progeny cells. These cells form, in case of *I. hospitalis*, the second membrane outside the cell cluster and the cell cycle re-starts with the formation of the cell division rings. In case of *Ca.* ‘I. morulus’, the outer membrane of the daughter cells is formed within the cell cluster. Rupture of the parental outer membrane releases progeny cells of *Ca.* ‘I. morulus’ with two membranes.

Since we were unable to observe inner compartments with double membranes inside an *I. hospitalis* cluster, we asked whether this type of cellular organisation might be more readily visible in other *Ignicoccus* species. Strikingly, in cultures of *Ca.* ‘I. morulus’ we observed cases in which a cluster was filled with cells that had distinct inner and outer membranes (Figures 4F and 4G), showing that the formation of the double membrane can be initiated before cells burst out of the parental outer membrane. Thus, despite being genetically closely related and sharing a common life cycle, *Ignicoccus* strains differ in the precise timing of outer membrane rupture of the cluster and double membrane re-generation in daughter cells (Figure 4H).

## Discussion

Compared to the biology of cell division in eukaryotes and bacteria, little is known about archaeal cell division. Most of what we know comes from the study of FtsZ-based division in *Haloferax* and CdvABC-based cell division in *Sulfolobus acidocaldarius*, respectively, both of which are ‘monoderms’ (*21*, *27*, *41–43*).

In this study of cell division in *Ignicoccus*, which is a diderm, we provide the first evidence that homologues of CdvA and the ESCRT-III proteins execute membrane constriction and scission in this archaeon. In line with recent data from *S. acidocaldarius* (*39*), our data suggest that *Ignicoccus* CdvA forms a polymeric division ring that constricts along with the ESCRT-III ring as cytokinesis proceeds. Remarkably through this analysis we also discovered that *Ignicoccus* strains appear unique to other archaea (*26*) in using polymeric rings to divide their inner membrane several times without deforming the outer membrane that separates the cell from its environment. This leads to the formation of cell clusters. Internal daughter cells then emerge later via rupture of the parental outer membrane in a manner that resembles the pop-out mechanism of reproduction in colonial algae like *Volvox* (*44*).

This is strikingly different from the way diderms, like *E. coli*, divide, which use physical connections between inner and outer membranes to deform and divide both membranes simultaneously in a way that ensures the preservation of cell structure over division cycles (*45*). In *E. coli*, this depends on a peptidoglycan cell wall, which *Ignicoccus* lacks. Instead, the cell division in *Ignicoccus* lineages might resemble cell division in the diderm bacterium *Thermotoga maritima* and related bacterial species, which have been reported to form clusters that are bounded by an outer membrane or “toga” (*46–50*). While the complete lifecycle and cell division of these bacteria remain poorly understood it is possible that some of them may have a life cycle similar to *Ignicoccus*.

Rupture of the outer membrane is not enough to complete the life cycle of *Ignicoccus* (Figure 4I). A key additional step in the restoration of the parental cell organisation is the regeneration of a double membrane. Strikingly, this appears to occur at different times in the different *Ignicoccus* species. In *Ca.* ‘I. morulus’ the double membrane of daughter cells emerged prior to rupture of the parental outer membrane, leading to the formation of cell clusters containing inner compartments that are effectively bounded by three membranes. Conversely, in *I. hospitalis* and *I. pacificus,* this process appears to occur concomitantly or soon after inner cells escape the bounding outer membrane. In these cultures, we frequently observed isolated cells with a single bounding membrane by imaging unfixed cells and by ExM and cryo-EM.

While future work will be required to define the precise mechanism of membrane duplication in *Ignicoccus*, we were able to catch cells in the act of double membrane formation by fluorescence-based ExM. In these intermediate states, the outer membrane appears to have ‘split’ locally to generate a region in which the two membranes are separated from one another by an intervening aqueous space – in a way that resembles a blister. This blister is separated from the rest of the single bounding membrane by two three-way membrane junctions. Intriguingly, when stained using antibodies against the outer membrane protein ACS, we were able to observe the protein partitioning into the outer membrane during the membrane duplication process (*40*). As a result, ACS is present in the membrane of cells having only one membrane. In addition, we found ACS protein in the outer membrane of cells with partially duplicated membranes and in the outer membrane of cells with distinct differentiated inner and outer membranes.

In summary, our exploration of the life cycle of *Ignicoccus* has revealed transitions in state as single cells give rise to clusters with multiple cytoplasms or daughter cells, as the cultures shift between the different growth phases. To the best of our knowledge, these different states have not been previously reported in archaea - emphasizing the importance of studying cell biology across different stages of a culture’s growth cycle and across archaeal strains.

The identification of *Ignicoccus* clusters that contain large numbers of daughter cells, which are related by division, helps to emphasize the fact that multicellular-like phases and different growth modes are not an exclusive feature of eukaryotes (*51*) and bacteria, like *Myxobacteriaceae* (*52*). Also the recent work on *Haloferax* cells under confinement showed multicellular-like cell architectures, but forced by mechanical compression (*53*). Strikingly, however, *Ignicoccus* is remarkable in changing its complex membrane architecture to cycle between states with one, two or three bounding membranes as it shifts from single cells to clusters and back again. We argue that this represents a new mode of cell division and cellular lifestyle.

Considering the diversity of cell architecture and distinct cell division protein repertoires among archaea, particularly the Asgard archaea (*15*, *43*, *54–57*), we envision that the further exploration of cytokinesis modes among different representatives across the archaeal tree of life will be crucial not only to better understand archaeal cell biology, but also to trace the origins of complex eukaryotic organization, and to understand how cells with multiple membranes preserve their form as they grow and divide.

## Author contributions

**F.M.** – experimental design; cloning, expression and purification of cell division proteins; antibody generation; large scale cultivation and cell processing; development of all membrane staining techniques; established staining of intermembrane compartment and DNA; imaging of non-fixed cells; established mounting of cells on agarose pads and Fluoro dishes; analysis of growth phase-dependent cell size parameters; established expansion microscopy for *Ignicoccus* strains; immunolocalization of cell division proteins in *I. hospitalis* and *I. pacificus*; preparation of samples for mass spectrometry; analyzed cell division ring diameter; cultivation of cells for Cryo-EM and Cryo-FIB-SEM; preliminary DNA isolation of all *Ignicoccus* strains and Oxford Nanopore sequencing; staining and analysis of cell cluster and progeny cells of all *Ignicoccus* strains; imaged pop-out mechanism of progeny cells; investigated membrane duplication process with ACS antibody; development of the life cycle model together with B.B.; analysis and interpretation of data, paper writing;

**R.S.-** developed an improved DNA isolation protocol for *Ca.* ‘Ignicoccus morulus’ and sequenced DNA using Oxford Nanopore sequencing technology; sequencing data analysis; species tree with all *Ignicoccus* strains.

**A.v.K., Z.W., T.A.M.B. –** FIB-SEM workflow development, data collection, analysis and manuscript editing

**J.P. -** introduction to staining and immunostaining techniques

**H.H.** – Isolation of *Ca.* ‘Ignicoccus morulus’ strain; large scale cultivation of *I. pacificus*, *I. islandicus* and *Ca.* ‘Ignicoccus morulus’

**R.R. and V.S. –** technical support in large scale cultivation of *I. hospitalis*

**D.G., A.S. and B.B. -** project conceptualisation, analysis and interpretation of data, paper writing, funding.

## Resource availability

Requests for further information and resources should be directed to and will be fulfilled by the corresponding authors.

## Materials availability

*Ignicoccus* strains are available at the German Collection of Microorganisms and Cell Cultures GmbH (DSMZ) under the DSM numbers. DSM 18386 for *Ignicoccus hospitalis*, DSM 13166 for *Ignicoccus pacificus* and DSM 13165 for *Ignicoccus islandicus* (DSM). Plasmids and antibodies generated in this study will be shared by the corresponding authors with a materials transfer agreement.

## Data and code availability

Any additional information required to reanalyze the data reported in this paper is available from the corresponding authors upon request.

## Acknowledgements

Mass spectrometry analysis was performed at the Biological Mass Spectrometry and Proteomics Facility of the Medical Research Council Laboratory of Molecular Biology, Cambridge, UK. We would like to thank Dr. Catarina Franco and Dr. Tom Smith for the help with data analysis and helpful discussions on experimental design.

electron microscopy facility for supporting sample preparation and data collection. Large scale cultivation was performed at the Archaea Centre of the University of Regensburg, Germany. We would like to thank Dipl. Ing. (FH) Thomas Hader, Gabi Gmeinwieser and Lena Kampf for help with large scale cultivation. We thank Dr. Felix Mikus and Dr. Gautam Dey (EMBL, Heidelberg, Germany) for help with expansion microscopy set-up. Further, we thank Laura Villanueva (NIOZ, Texel, Netherlands) for joint efforts to secure funding and thank all group members of the Grohmann, the Baum and the Spang lab for help and fruitful discussions.

We thank the Life Sciences-Moore-Simons foundation (735929LPI) and a Gordon and Betty Moore Foundation’s Symbiosis in Aquatic Systems Initiative (GBMF9346) for supporting our collaborative work together.

Furthermore, A.S. has received funding from the European Research Council (ERC) under the European Union’s Horizon 2020 research and innovation programme (grant agreement No. 947317, ASymbEL).

B.B. received support for this work from the Medical Research Council - Laboratory of Molecular Biology (MC_UP_1201/27) and the Wellcome Trust (222460/Z/21/Z).

D.G. thanks the University of Regensburg for core funding of the German Archaea Centre and gratefully acknowledges funding by Deutsche Forschungsgemeinschaft in the German-Israeli Project Cooperation Programme (EL 1199/2-1 | SCHW 716/15-1).

T.A.M.B.’s laboratory is supported by the Medical Research Council, as part of UK Research and Innovation (Programme MC_UP_1201/31). T.A.M.B. thanks the Wellcome Trust (grant 225317/Z/22/Z).

## Methods

### Cloning of cell division genes

Cell division genes of *Ignicoccus hospitalis* DSM 18386 were cloned into the vector pET-19b (Novagen, Sigma-Aldrich No. 69677), containing and N-terminal Histidin (10)-tag, an IPTG inducible promoter and an ampicillin resistance cassette. The genes Igni_0996 (CdvA), Igni_0995 (CdvB), Igni_1156 (CdvB1) and its corresponding pET-19b backbones were amplified by gradient-PCR using specifically designed starter oligonucleotides (Table S1). 50 ng genomic DNA of *I. hospitalis* or 10 ng plasmid DNA was used as template DNA. PCR products were amplified with Phusion High-Fidelity DNA Polymerase (Thermo Fisher Scientific, No. F530S) using a temperature gradient from 55 to 65°C. PCR products were separated on 1% (w/v) agarose gels (Molecular Biology Grade Agarose, Eurogentec, No. EP-0010-10) and stained in a 0.5 µg/mL ethidium bromide solution. Samples containing PCR products with correct size were pooled and template DNA was digested with the restriction enzyme DpnI (New England Biolabs, No. R0176S) over night at 37°C. PCR products were purified using the Qiagen QIAquick PCR Purification Kit and eluted in 30 µL sterile deionized water. The DNA concentration was determined using absorption spectroscopy employing a NanoDrop ND-1000 Spectrophotometer (Thermo Scientific). Gibson Assembly was used to clone the genes into pET-19b. Specifically, a self-made Gibson Assembly mix (in total 200 µL, split into 15 µL reaction portions and stored at −20°C) was used containing 53.33 µL 5x ISO-buffer, 0.11 µL T5 Exonuclease (10 U/µL, New England Biolabs, No. M0663S), 26.67 µL Taq DNA ligase (40 U/µL, New England Biolabs, No. M0208S), 3.33 µL Phusion High-Fidelity DNA Polymerase (2 U/µL, Thermo Fisher Scientific, No. F530S) and 116.56 µL H2O. The composition of the 5x ISO-buffer (for 2 mL) was, 1 mL Tris-HCl (1 M, pH 7.5), 100 µL MgCl2 (1 M), 20 µL dATP (0.1 M), 20 µL dGTP (0.1 M), 20 µL dTTP (0.1 M), 20 µL dCTP (0.1 M), 100 µL Dithiothreitol (1 M), 0.5 g PEG 8000, 100 µL NAD (0.1 M), ad 2 mL with H2O. The 5x ISO-buffer was stored at −20°C. For Gibson Assembly, 5 µL of DNA (1:4 ratio, plasmid:insert) was added to 15 µL Gibson Assembly mix and assembly was performed for 1 h at 50°C in a thermocycler. The complete reaction (20 µL) was used for heat-shock transformation (42°C, 75 sec) of chemical-competent *E. coli* DH5α cells (ThermoFisher Scientific, No. 18265017) and positive clones were selected on LB plates containing 100 µg/mL ampicillin. Clones were picked and cultivated in 5 mL LB medium for 15 h at 37°C and under shaking of 225 revolutions per minute (rpm) in an incubator (New Brunswick Innova 44). Cells were sedimented at 10,000 x *g* for 1 min and plasmids were isolated using the peqGold Plasmid Miniprep Kit II (VWR). Plasmid DNA was eluted in 30 µL sterile deionized water. To screen for possible positive clones, containing the correct insert, plasmids were cut once with the restriction enzyme EcoRI-HF (New England Biolabs, No. R3101S) to linearize the plasmids and the size of the linearized plasmids was estimated on an 1% (w/v) agarose gel. Plasmids with the correct size were further analyzed by Sanger sequencing (BaseClear, Leiden, NL) using the primer T7 (5’ to 3’ sequence: TAATACGACTCACTATAGGG) and/or T7 Term primer (5’ to 3’ sequence: TGCTAGTTATTGCTCAGCGG) to sequence the cloned insert including the His-Tag region. The successfully cloned plasmids were named pFMA2 for Igni_0996 (CdvA), pFMA3 for Igni_1156 (CdvB1) and pFMA9 for Igni_0995 (CdvB), respectively.

### Recombinant expression and purification of cell division proteins

For recombinant expression of cell division proteins, the vectors pFMA2, pFMA3 and pFMA9 were transformed by heat-shock (42°C, 75 sec) into *E. coli* Rosetta (DE3) pLysS (Novagen, Sigma-Aldrich No. 70956), which contains the pRARE1 plasmid. The plasmid pRARE1 contains a chloramphenicol-resistant cassette and supplies the tRNAs for unusual codons (AGG, AGA, AUA, CUA, CCC, GGA) rarely used in *E. coli*. This strain was used for recombinant expression of Cdv proteins because all cell division genes show codons encoding for unusual tRNAs which are not present in strains like *E. coli* BL21(DE3). Positive transformants containing the cloned plasmids and the pRARE plasmid were selected on LB plates containing 100 µg/mL ampicillin and 34 µg/mL chloramphenicol. Transformed *E. coli* cells were cultivated in 1.5 L Erlenmeyer flasks containing 250 mL LB medium with ampicillin (100 µg/mL) and chloramphenicol (34 µg/mL). The medium was inoculated with an overnight grown pre-culture to an OD600 between 0.2 and 0.3, and was incubated at 37°C under shaking at 225 rpm. Optical density was measured every hour, and protein expression was induced by the addition of 1 mM IPTG after reaching an OD600 of 1.0. Before induction and every hour after induction (for 4 h long), 1 mL samples were taken, cells were sedimented by centrifugation and used for SDS-PAGE to monitor protein expression. 4 hours after induction, all cells were sedimented by centrifugation at 5,000 x g for 10 min and 4°C (Eppendorf, Centrifuge 5804R). For protein purification, cells were lysed with B-PER Bacterial Protein Extraction Reagent (ThermoFisher Scientific, No. 90084) with addition of DNase I (Roche, No. 10104159001). Cell debris was removed by centrifugation for 15 min at 5,000 g and the supernatant was separated in cytoplasm and membranes by ultra-centrifugation for 2 h at 45,000 rpm. To purify the proteins via the N-terminal Histidine (10)-tag, Ni-NTA agarose was equilibrated with an equilibration buffer containing 50 mM NaH2PO4, 300 mM NaCl, 10 mM imidazole (pH 7.5). The cytoplasmic fraction was incubated with the equilibrated Ni-NTA agarose (2 mL) for 30 min under gently shaking at 4°C. Ni-NTA agarose was transferred to Protino Columns (Machery-Nagel, No. 745250.10), washed twice with wash buffer (50 mM NaH2PO4, 300 mM NaCl, 20 mM imidazole, pH 7.5) and the protein was eluted with an imidazole gradient by adding stepwise elution buffer (50 mM NaH2PO4, 300 mM NaCl, x mM imidazole, pH 7.5) with different imidazole concentrations (x = 70, 90, 110, 130, 150 mM), followed by a final addition of elution buffer containing 800 mM imidazole. The successful protein purification was monitored by SDS-PAGE using 12% Mini-PROTEAN TGX Precast Protein Gels, (12-well, 20 µl, Biorad) and separated proteins stained with Coomassie.

### Raising of polyclonal antibodies against cell division proteins in *I. hospitalis*

Polyclonal antibodies were raised in different hosts by Davids Biotechnologie (Regensburg, Germany). For immunization, antigen was extracted from SDS gel pieces according to manufacturer’s protocol. The antigen CdvA (Igni_0996) was used to immunize chicken. The hen was immunized with protein 4 times in a period of 35 days, followed by a first egg collection on day 50. Afterwards the hen was once again immunized with CdvA protein. The final egg collection took place on day 63 and the antibody was prepared from the egg yolk according to manufacturer’s protocols. The antibody for CdvB (Igni_0995) was produced in a rabbit, which was immunized 5 times with antigen over a period of 56 days with antiserum collection on day 63. The antibody for CdvB1 (Igni_1156) was raised in two guinea pigs, which were both immunized 5 times with antigen within 56 days and afterwards the antiserum was collected on day 63.

### Cultivation of *Ignicoccus hospitalis* in a 300 L bioreactor

*Ignicoccus hospitalis* KIN4/I (DSM 18386, DSMZ Braunschweig, Germany) was cultivated over a time period of 24 h at 90°C in a 300 Liter bioreactor with 280 Liter ½-SME medium (*58*), 1 kg sulphur, 280 g of yeast extract and under a continuous gas flow of 15 L/min H2/CO2 (80/20, v/v) under a pressure of 2 bars. The pre-cultures for bioreactor inoculation (12x 1L-pressure bottles, each with 250 mL ½ SME medium with 5 g/L sulphur and H2/CO2 (80/20, v/v, 1 bar) were cultivated for 19 h at 90°C and under shaking (35 rpm). The bioreactor was inoculated with liquid pre-culture and a stationary-phase cell pellet (2 g) from a former bioreactor cultivation. Bioreactor samples, each with a volume of 30 L, were taken 4, 6, 8 h after inoculation and were immediately cooled down. Remaining sulphur settled over night and cells in the supernatant were harvested with a flow-through centrifuge at 165 mL/min, 4°C and 15,500 rpm (Heraeus Contifuge Stratos). The remaining cells were sedimented by centrifugation at 5,000 x *g*, 4°C for 30 min (JA-10 rotor, Beckman Coulter Avanti J-25), frozen with liquid nitrogen and stored at −70°C. After cultivation of *I. hospitalis* for 24 h the bioreactor was cooled down and the remaining 190 L cell broth was harvested over night with a Padberg centrifuge at 20,000 rpm with a flow rate of 200 mL/min at 4°C. Afterwards, the cells were scratched from the Teflon membrane and resuspended in the remaining rotor liquid. Again, a sample was taken for electron microscopy and stored at 4°C. The remaining cells were sedimented by centrifugation at 5,000 x *g*, 4°C for 30 min (JA-10 rotor, Beckman Coulter Avanti J-25), frozen with liquid nitrogen and stored at −70°C.

### Cell viability test of *Ignicoccus* cells in a state of suspended animation

To examine the cell viability of *Ignicoccus* cells, which are in a state of suspended animation due to freezing in liquid nitrogen and storage at −70°C, a small amount of cells, broken apart from a frozen cell pellet, were resuspended in 200 µL of ½ SME medium. A volume of 100 µL cell suspension was used to inoculate septum flasks with 20 mL of ½ SME medium with sulphur (5 g/L) and H2/CO2 (80/20, v/v, 2 bar) gas phase. Cultivation was performed at 90°C under shaking at 70 rpm. The total cell number was determined by cell counting directly after inoculation of the medium (t = 0) and finally after 24 h of cultivation at 90°C (t = 24). The experiment was performed using cell material from frozen samples taken at time points 4 h, 6 h, 8 h and 24 h during 280 L-bioreactor cultivations (N = 3).

### Cultivation of Ca. ‘Ignicoccus morulus’, Ignicoccus islandicus and Ignicoccus pacificus

*Ca.* ‘Ignicoccus morulus’ (Mex13A-SI-L1A) was cultivated over a period of 30 h at 90°C in a bioreactor with 250 L of ½-SME medium with 1 kg of sulphur and without yeast extract (pH 6.0). Initially, cells were cultivated for 7.5 h with H2/CO2 (80/20, v/v) at 2 bar, followed by cultivation with a continuous gas flow of 55 L/min with N2/H2/CO2 (65/15/20, v/v/v) for 22.5 h at 2 bar. The bioreactor was inoculated with 12x 1L-pressure bottles, each with 250 mL ½-SME medium with 5 g/L sulphur and H2/CO2 (80/20, v/v, 1 bar). Pre-cultures were cultivated at 90°C and under shaking (35 rpm) until a cell density of 2 × 107 cells/mL was reached. After bioreactor cultivation cells of *Ca.* ‘Ignicoccus morulus’ were harvested with a Padberg centrifuge at 20.000 rpm with a flow rate of 200 mL/min at 4°C. Cultivation of *Ignicoccus islandicus* and *Ignicoccus pacificus* was performed in 250 L of ½-SME medium with 1 kg of sulphur and 0.1% (w/v) yeast extract and cultivation was carried outin the same way as described for *Ca.* ‘Ignicoccus morulus’, with the following exceptions: For *I. islandicus*, cultivation time in the bioreactor was 23 h, first for 8.5 h with H2/CO2 (80/20, v/v, 2 bar) followed by cultivation with a gas flow of 45 L/min with N2/H2/CO2 (65/15/20, v/v/v). Cells were harvested after the cell density reached 6.3 × 107 cells/mL. For *I. pacificus*, cultivation time was 26 h in the bioreactor. Initially for 7.85 h of cultivation with H2/CO2 (80/20, v/v, 2 bar) followed by a gas flow of 55 L/min with N2/H2/CO2 (65/15/20, v/v/v). Cells were harvested after reaching a cell density of 1 × 108 cells/mL. Precultures for *I. islandicus* and *I. pacificus* were cultivated for 16 h at 90°C under shaking.

### Membrane staining of unfixed *I. hospitalis* cells

For staining of *I. hospitalis* cells with different membrane dyes, the cells were first washed twice with PBS and sedimented by centrifugation at 5,000 x g for 3 min at 4°C. Dyes were diluted in PBS and cells resuspended in 100 µL dye-PBS solution and stained under shaking (300 rpm). For ConA-647 (C21421, Invitrogen) staining, the dye was diluted 1:40 and cells were stained for 2 h. For CellTracker Red CMPTX (C34552, Invitrogen) staining, the dye was diluted 1:5,000 and cells were stained for 1 h. For CellMask Deep Red (C10046, Invitrogen) staining, the dye was diluted 1:10,000 and cells were stained for 30 minutes. For multicolor staining of *I. hospitalis*, cells were first stained with CellMask Deep Red for 30 min, washed three times with PBS and afterwards stained for 1 h with ConA-488 (C11252, Invitrogen) diluted 1:40 with PBS. For Bodipy TR Ceramide (D7540, Invitrogen) staining, the dye was diluted 1:2,000 and cells were stained for 1 h. In all cases, cells were sedimented after staining by centrifugation at 5,000 x *g* for 3 min at 4°C and were washed three times with PBS buffer. Finally, cells were resuspended in DNA staining solution, containing DAPI, which was 1:1,000 diluted with PBS. Cells were immobilized on agarose pads as described in the corresponding methods section.

### Dextran staining of *I. hospitalis*

To investigate membrane permeability in *I. hospitalis*, Oregon Green 488-labeled dextrans of a molecular mass of 10 kDa (D7170, Invitrogen) or 70 kDa (D7173, Invitrogen) were used. Before staining, cells were washed twice with PBS and sedimented by centrifugation at 5,000 x *g* for 3 min. Cells were resuspended in dextran solution (final conc. 10 µg/mL) and incubated at 300 rpm on a shaker allowing passive diffusion of dextran into the cell for 1 h at room temperature. Afterwards cells were sedimented by centrifugation at 5,000 x *g* for 3 min at 4°C and were washed three times with PBS. Cells were resuspended in DAPI solution (1:1,000 in PBS) and were immobilized on agarose pads as described in the corresponding methods section.

### Membrane staining of unfixed cells from *I. islandicus*, *I. pacificus* and *Ca.* ‘I. morulus’

To investigate the number of progeny cells in *I. islandicus*, *I. pacificus* and *Ca.* ‘I. morulus’, the same dyes CellTracker Red CMTPX (C34552, Invitrogen) and MitoTracker Green FM (M7514, Invitrogen) were used but different methods to adhere the cells. This is due to the different membrane characteristics that differ between the strains. For example, not all *Ignicoccus* strains have glucose- and/or mannose glycosylated lipids and therefore these strains do not bind to ConA-coated dishes. In all cases, cells were first washed twice with PBS, sedimented by centrifugation at 5,000 x *g* for 3 min at 4°C and were resuspended in PBS or dye-PBS solution, depending on the cell immobilization method. In detail *I. islandicus* was immobilized on poly-L-lysine coated Fluoro dishes. To this end, FluoroDish cell culture dishes (35 mm, 23 mm glass bottom, No-FD35-100, WPI) were coated for 30 min with 0.1% (w/v) poly-L-lysine solution (P8920, Sigma-Aldrich) and washed three times with PBS. The cell pellet of *I. islandicus* was resuspended in 200 µL PBS and dropped onto the poly-L-lysine Fluoro dish and incubated at room temperature for 30 min without shaking. After that, the liquid was removed and the adhered cells were washed three times with PBS. 200 µL of dye-PBS solution, containing CellTracker Red CMTPX (1:10,000) and Mitotracker Green FM (1:10,000) was dropped onto the Fluoro dishes and incubated for 1 h on a platform rocker (Polymax 2040, Heidolph) at 25 rpm. Cells were washed three times with 500 µL PBS and then 200 µL of PBS-DAPI (1:1,000, DAPI stock solution 1 mg/mL) was dropped onto the cells and imaged with the confocal spinning disc microscope. *Ca.* ‘I. morulus’ cells were stained and immobilized on Fluoro dishes as described for *I. islandicus*, but adhered with ConA (0.5 mg/mL dissolved in water) instead of poly-L-lysine. For *I. pacificus* cells, agarose pads were used. In detail, the washed cell pellet was resuspended in dye-PBS solution (CellTracker Red CMTPX (1:10,000) and MitoTracker Green FM (1:10,000)) and stained for 1 h at room temperature on a shaker at 300 rpm. Cells were sedimented by centrifugation at 5,000 x *g* for 3 min at room temperature and were resuspended in DAPI-PBS (1:1,000, DAPI stock solution 1 mg/mL) and 10 µL of stained *I. pacificus* cells were dropped onto agarose pads as described earlier and imaged.

### Immobilization of cells on agarose pads

1% (w/v) low-melt agarose (Sigma Aldrich, A9414-100G) was dissolved in autoclaved MilliQ-water by heating the solution in a microwave. 200 µL molten agarose was dropped onto a microscope slide containing a spacer, each with 4 layers tape, one both sides of the slide. The agarose drop was squeezed to a pad with a second microscope slide on top and polymerized for 10 min. Afterwards the top microscope slide was removed and 10 µL fluorescence-labeled sample was applied. Immediately, the solution was spread on the whole agarose pad by gently agitation. After the surface of the pad dried completely a round cover glass (13 mm in diameter, thickness no. 1.5, VWR, 631-0150) was placed on top of the agarose pad and the sample was imaged.

### Expansion Microscopy and immunofluorescence staining

Cells of *Ignicoccus* were washed twice with PBS, sedimented by centrifugation for 3 min (5,000 x g, 4°C) and fixed for 20 min with 3.7% (w/v) formaldehyde (in PBS) under shaking (400 rpm) at room temperature. Cells were sedimented again and washed twice with 1 mL PBS. The cell pellet was resuspended in 200 µL AA/FA solution (1% acrylamide (w/v) + 0.7% (w/v) formaldehyde diluted in PBS) and incubated overnight (20°C, 400 rpm). Glass cover slips (6 mm) were coated with 35 µL Poly-L-Lysine solution (0.1% (w/v), Sigma-Aldrich P8920) for 30 min at room temperature. In parallel, a glass petri dish was lined with parafilm and cooled down on ice. The cover slips were washed three times with PBS and then 35 µL cross-linked cells were dropped onto it. Cells were adhered for 30 min to the cover slips and then the liquid was removed. Quickly, 1 µL of APS (10% (w/v) in H2O) and 1 µL TEMED (10% (v/v) in H2O) were added to an 18 µL monomer solution (19% (w/v) sodium acrylate, 10% (w/v) acrylamide, 0.1% (w/v) N,N’-methylenbisacrylamide; in 10x PBS). Upon mixing, two drops of each 9 µL were pipetted onto the glass petri dish lined with parafilm. The cover slips with the cells were placed onto the drops and incubated on ice for 5 min. Then the petri dish was transferred into a closed box containing wet paper towels to ensure a humid atmosphere and incubated for 1h at 37°C for gelation. The gels, attached to the cover slips, were incubated for 1h in denaturation buffer (8 M urea, 25 mM EDTA, 10% (w/v) SDS, pH 6.5-7.0) at 95°C. Subsequently, the cover slip was removed and the gel was washed three times with PBS. To permeabilize and expand the cells, the gels were incubated in Millipore H2O in a 6-well plate under shaking for 30 min. The water was replaced twice and the gels were stored overnight in H2O at 4°C, allowing full gel expansion. After cutting the expanded gels in pieces, e.g. quarter, long-time storage was done in PBS with 0.02% (w/v) sodium azide at 4°C after shrinking the gels in PBS. For immunostaining of cells with antibodies, the gel pieces were washed 3 times with PBS under shaking (400 rpm) first and then incubated in primary antibody solution (antibody 1:1,000 in PBS with 0.1% (v/v) Tween-20 and 3% (w/v) BSA) overnight at 4°C under shaking (400 rpm). Gel pieces were washed three times with PBS-T (PBS and 0.1% (v/v) Tween-20) for 10 min under shaking and incubated afterwards for 2 h in secondary antibody solution (secondary antibody 1:1,000 in PBS-T and 3% (w/v) BSA) and membrane dyes, e.g. Concanavalin A (ConA) conjugated with Alexa Fluor 647 (Invitrogen C21421, 1:40 dilution) or Bodipy TR Ceramide (Invitrogen D7540, 1:2,000 dilution) under shaking at room temperature and protected from light. Gel pieces were washed again three times with PBS and were then re-expanded in Millipore H2O in a 6-well plate (3 changes of water, incubation each 10 min). The expanded gel was placed on a with Poly-L-lysine solution coated cover slip (25 mm diameter, thickness no. 1, VWR No.631-0171) in the correct orientation (side with cells towards the cover slip) and cells were imaged by Spinning Disk Confocal Microscopy as described in the corresponding methods section.

### Imaging by Spinning Disk Confocal Microscopy

Imaging was performed on an inverted microscope (Nikon Eclipse Ti2) equipped with a SoRa scanner unit (Yokogawa) for confocal microscopes and a Prime 95B back-illuminated sCMOS (scientific Complementary Metal-Oxide Semiconductor) camera (Teledyne Photometrics). Images were acquired using an oil immersion objective (Apo TIRF 100x/1.49, Nikon) with a Low Autofluorescence Immersion Oil Type-F (Immoil-F30CC, Olympus). Using the 2.8x magnification lens in the SoRa scanner unit a total magnification of 280x could be achieved. Images were acquired with a 500 ms exposure time for the DNA stain DAPI and a 200 ms exposure time for fluorescence-labeled proteins or lipids. Image analysis was performed with the program Fiji.

### Cryo-FIB-SEM volume imaging and sample preparation

For cryo-FIB-SEM (cryogenic-Focused Ion Beam-Scanning Electron Microscopy) sample preparation, *Ignicoccus* cell culture was vitrified with a High Pressure Freezing machine (HPF) Compact 03 (Wohlwend, GmbH) using the previously described Waffle method (*59*). For each sample, two type-A gold planchettes (3 mm) were coated with 0.1 % (w/v) lecithin dissolved in chloroform, and one 200 mesh Au-continuous carbon grid was glow discharged for 30 sec at 40 mA. 2.5 µL of dense cell culture was applied to the grid-bar side of the EM grid and sandwiched between the two HPF planchettes before jet-freezing in the high-pressure freezer. Frozen grid-planchette assemblies were dissembled under liquid nitrogen and grids were clipped into auto-grid rings (ThermoFisher). Cryogenic volume SEM was performed as previously described by Schertel (*60*) with the following modification: cryo-FIB-SEM data were acquired with a Crossbeam-550 (Zeiss) with a FIB column positioned 54° relative to the SEM column. SEM images were acquired at a lateral pixel-size of 5.7 nm (*I. hospitalis*) or 4.4 nm (*I. pacificus*) at an acceleration voltage of 2.33 kV with 41 pA probe using an in-lens detector. After each SEM acquisition a 20-nm slice was removed using FIB operating at 30 kV with a 300 pA probe. This imaging-slicing cycle iterated to produce the volume EM data. Raw-image stacks were drift-corrected using linear stack alignment with Scale-Invariant Feature Transform (SIFT) (*61*) algorithm implemented in Fiji (*62*). Curtaining and charging artefacts were afterwards removed using cryo-FIB-SEM Image stacks tool (https://github.com/EMCRUMC/napari-cryofibsem-imgproc) implemented in napari (https://zenodo.org/records/15779115) using default settings. Images were scaled by 1.236 (1/sin(54°)) in Y to compensate for the tilted SEM view. Final images were Fourier band-pass filtered in Fiji using default settings.

### Cryo-ET and sample preparation

For cryo-EM grid preparation, the culture was mixed 1:10 with 10 nm protein-A gold (CMC Utrecht, Netherlands) and then 2.5 µL of the culture was applied to a freshly glow-discharged Quantifoil R3.5/1 Au 200 mesh grid, adsorbed for 2 s, blotted for 5 s and plunge-frozen into liquid ethane using a Leica EM GP2. Cryo-ET (Electron Tomography) data was collected on a Titan Krios G3 microscope equipped with a Quantum energy filter (slit width 20 eV) and a K3 direct electron detector running in counting mode with SerialEM (*63*). Tilt series were collected with a defocus range of −5 to −8 µm, between ±60° in a grouped dose symmetric scheme (*64*) with a 2° tilt increment. A total dose of 175 e-/Å2 was applied over the entire series, and image data were sampled at a pixel size of 2.809 Å with 10 fractioned frames per tilt image. Unaligned tilt-movie frames were motion-corrected in IMOD (*65*) and tilt series alignment using gold fiducials was performed in IMOD. Tomograms for visualisation were generated using IMOD and denoised with Cryo-CARE (*66*). Figure panels containing cryo-ET images were prepared using Fiji (*62*).

### Proteomic analysis

*I. hospitalis* cell pellets were suspended in 500 µL of 50 mM ammonium bicarbonate (AMBIC) and sonicated on an icy bath at an amplitude of 28% for a total of 90 sec with 10 sec on/10 sec off cycles. RapiGest (Waters) was then added to the cell lysates to a final concentration of 0.05% (v/v). For mass spectrometry analysis, 50 µg of each cell lysate sample was diluted to approximately 0.6 µg/µL using 25 mM AMBIC. Cysteines were reduced by adding DTT to a final concentration of 4 mM and heating the samples to 60°C for a total of 10 min. To prevent cysteine re-oxidation, iodoacetamide was added as an alkylating reagent to a final concentration of 14 mM and incubation proceeded for 45 min at room temperature in the dark. Digestion was carried out semi-automatically on a Kingfisher Apex using an adapted Protein Aggregation Capture method (*67*, *68*). Briefly, reduced and alkylated samples were transferred to a 96-well plate and precipitated by adding acetonitrile to a final concentration of 70% (v/v). Washed MagResyn Hydroxyl microparticles (Resyn Biosciences) were immediately added to the samples at a ratio of 1:4 (protein:bead) to promote protein precipitation and on-bead aggregation. Three subsequent washes of the beads with the aggregated proteins was performed with 100% acetonitrile and followed by two additional washes with 70% (v/v) ethanol. In-bead digestion was performed on a first stage on the Kingfisher Apex by adding 1 µg of trypsin to 100 µL of 25 mM AMBIC containing 0.2% RapiGest detergent and incubating for 1 h at 47°C. Overnight digestion was then carried out at 37°C on an Eppendorf ThermoMixer C for an additional 16 h. Magnetic beads were removed and peptides were acidified with the addition of trifluoroacetic acid to a final concentration of 0.5% (v/v). The acidified tryptic digest was then centrifuged at 13,000 x g for 15 minutes to remove RapiGest degradation by-products and the supernatant subjected to LC-MS/MS analysis. LC-MS/MS was performed on an U3000 Ultimate HPLC (ThermoFisher Scientific, San Jose, USA) hyphenated to an Q-Exactive plus mass spectrometer (ThermoFisher Scientific, San Jose, USA). Peptides were trapped on a C18 Acclaim PepMap 100 (5 µm, 300 µm x 5mm) trap column (ThermoFisher Scientific, San Jose, USA) and separated on a C18 Aurora Ultimate TS (25cm x 75 µm) column (IonOpticks, Australia) over a gradient of solvent B [80 % (v/v) acetonitrile, 0.1% (v/v) formic acid] from 9% to 34% B over 110 min followed by a 95% B over 30 min before column equilibration in preparation for next sample injection. MS1 full scans were acquired in the Orbitrap at a resolution of 70,000 (AGC target of 1e5 ions with a maximum injection time of 100 ms) and followed by MS2 in a data-dependent acquisition setting composed of 15 TOP N with a minimum AGC target of 1e3 ions, at a resolution of 35,000 with a maximum injection time of 100 ms and an HCD collision energy of 27%. Raw data were imported and processed in Proteome Discoverer (v3.1) (ThermoFisher Scientific). Raw files were searched against *Ignicoccus hospitalis* protein sequences downloaded from UniProt in June of 2025 (UP000000262_45391).

### Proteomics data analysis and statistics

Peptide-level output from Proteome Discoverer was processed in R (v4.5.2) using QFeatures (v1.20.0, https://rformassspectrometry.github.io/QFeatures) and biomasslmb (v0.0.5, https://lmb-mass-spec-compbio.github.io/biomasslmb/) R packages. Peptides were filtered to remove matches to contaminants, log2-transformed and median normalized. Peptides with fewer than 3 quantification values were removed, before summarisation to protein-level abundances using the robustSummary function from MsCoreUtils (*68*). In total, 1,223 proteins out of the 1,434 proteins in the *Ignicoccus hospitalis* reference proteome (UP000000262) were quantified. Across the 12 samples, 355 values were missing (2.4%) and were imputed using the MinProb method implemented in QFeatures. Statistical testing was performed with limma (v3.66.0) (*69*) to compare protein abundance between timepoints. The treat function was used to set the null hypothesis as log◻ fold-change < 1.5, with a mean–variance trend fitted to the prior variance. Pairwise contrasts were computed between sequential timepoints. *P*-values were adjusted for multiple testing using the Benjamini-Hochberg false discovery rate (FDR) procedure (*70*) with proteins below an adjusted *p*-value of 0.05 considered statistically significant.

### DNA extraction and genome sequencing

Cells (0.1 g) of *Ca.* “Ignicoccus morulus” (Mex13A-SI-L1A), cultivated in a 300 L bioreactor as described above, were resuspended in 510 μL TE buffer (10 mM Tris, 1 mM EDTA, pH 8.0) and incubated at 56°C for 1 h with 15 μL Proteinase K (20 mg/mL), 5 μL RNase A (20 mg/mL) and 15 μL of 20 % SDS. NaCl was added to a final concentration of 0.3 M. The lysate was extracted twice with phenol:chloroform:isoamyl alcohol (25:24:1, (v/v/v), pH 8.0), followed by two extractions with chloroform:isoamyl alcohol (24:1, (v/v)). Phase separation was achieved via centrifugation at 21,500 x g for 2 min at room temperature. Genomic DNA was precipitated by adding 0.6 volumes of cold 100 % propan-2-ol and 0.06 volumes of 3 M sodium acetate (pH 5.2) and overnight incubation at −20°C. After centrifugation at 16,000 x g for 10 min at room temperature, precipitated genomic DNA was washed twice with icecold 70 % ethanol and after drying, the pellet was resuspended in 50 μL EB buffer (Qiagen). Size selection of the genomic DNA was performed as described earlier (https://dx.doi.org/10.17504/protocols.io.bwkdpcs6). Briefly, 30 μg of extracted DNA in EB buffer was diluted to a volume of 60 µL with 10 mM Tris-HCl (pH 8.0), mixed with an equal volume of size selection solution A (4 % (w/v) polyvinylpyrrolidone (MW 360,000), 1.2 M KCl, 20 mM Tris-HCl, pH 8.0) and centrifuged at 10,000 x g for 30 min at room temperature. The supernatant was centrifuged a second time and the pellets from both centrifugation steps were combined. The combined pellet was washed twice using 200 μl of 70 % (v/v) ethanol, sedimented by centrifugation at 10,000 x g for 3 min at room temperature and then resuspended in 30 μl EB buffer. Library preparation was done using the ONT Native Barcoding Kit 96 V14 (SQK-NBD114.96, Oxford Nanopore Technologies), following manufacturer’s instructions (NBE_9171_v114_revR_30Jan2025). Sequencing was performed for 96 h using a R10.4.1 flow cell (FLO-MIN114) on a MinION Mk1D (MinKNOW v25.03.9). Reads were base called using Dorado v1.0.0 (https://github.com/nanoporetech/dorado/releases/tag/v1.0.0) with the dna_r10.4.1_e8.2_400bps_sup@v5.2.0 model in simplex mode, resulting in 183 440 raw reads. The genome was assembled using Hybracter (v0.11.2) (*71*) [hybracter long –auto], resulting in a closed genome with a length of 1.54 Mb with a minimum coverage of 130.8, and an estimated CheckM2 (v1.1.0) (*72*) completeness of 99.64% and contamination of 0.25%.

### Phylogenetic analyses - CdvB phylogeny (SNF7 domain proteins)

SNF7 domain proteins of eukaryotes and Asgard archaea, i.e. homologs of Vps2-24-46 and Vps20-32-60, respectively (*15*) as well as of other archaea (i.e. proteins assigned to arCOG00454, arCOG00453 and arCOG00452 for a selected set of our in-house archaeal reference genomes including *Ignicoccus hospitalis* (see (*73*)) were concatenated and duplicates (i.e. sequences with 100% identity) removed. Sequences were aligned using MAFFT L-INS-I (v7.407) (*74*) trimmed with BMGE (v1.12) (*75*) [-m BLOSUM30 -b 2 -h 0.55] and subjected to phylogenetic analyses using IQ-TREE 2 (v. 2.1.1) (*76*) [-m LG+C20+R+F -bb 1000 -alrt 1000]. The tree was viewed in FigTree (https://github.com/rambaut/figtree/) and annotated using Adobe Illustrator. Protein structure models for CdvB, CdvB1 and CdvB2 of *I. hospitalis* were generated by AlphaFold 3 (https://alphafoldserver.com/) in December 2025 and superimposition was done with default settings in UCSF-ChimeraX (v1.10.1) (https://www.rbvi.ucsf.edu/chimerax/docs/credits.html).

### Species phylogeny

Maximum likelihood phylogenetic reconstruction of an archaeal species tree was performed using assemblies from the Genome Taxonomy Database (GTDB) (*77*). Briefly, the GTDB (r226, 16th April, 2025) was queried for ‘((“GTDB Taxonomy” CONTAINS “o Sulfolobales” AND “GTDB Representative of Species” IS TRUE AND “Assembly Level” IS NOT “contig”) OR (“GTDB Taxonomy” CONTAINS “o Nitrososphaerales” AND “GTDB Representative of Species” IS TRUE AND “Assembly Level” IS “complete genome”)) ‘. This resulted in 145 genomes from the order *Sulfolobales* and 36 genomes from the order *Nitrososphaerales*, which were used as an outgroup. Retrieved assemblies and the newly sequenced genome of *Ca.* “I. morulus” (Mex13A-SI-L1A) were searched for 53 archaeal genes based on a subset of the “top-ranked marker proteins” from a recent evaluation based on minimizing horizontal gene transfer and optimising the recovery of monophyletic lineages (*73*) via GTDB-Tk identify (v2.6.1) (*78*). The protein sequences of the detected marker genes were individually aligned using MAFFT L-INS-I (v7.525) (*79*) [--localpair -reorder --maxiterate 1000] and trimmed using BMGE (v1.12) (*75*) [-t AA -m BLOSUM30 -h 0.55]. A phylogenetic tree was then generated using IQ-TREE (*80*) (v3.0.1) [-m LG+C40+F+G -bb 1000 -alrt 1000] from the concatenated marker alignments [iqtree -p --out-aln --out-format FASTA]. The final tree was rooted with the outgroup and annotated using ape (*81*) (v5.8.1), ggtree (*82*) (v4.0.4) and treeio (*83*) (v1.34.0)

---

**Table S1:**
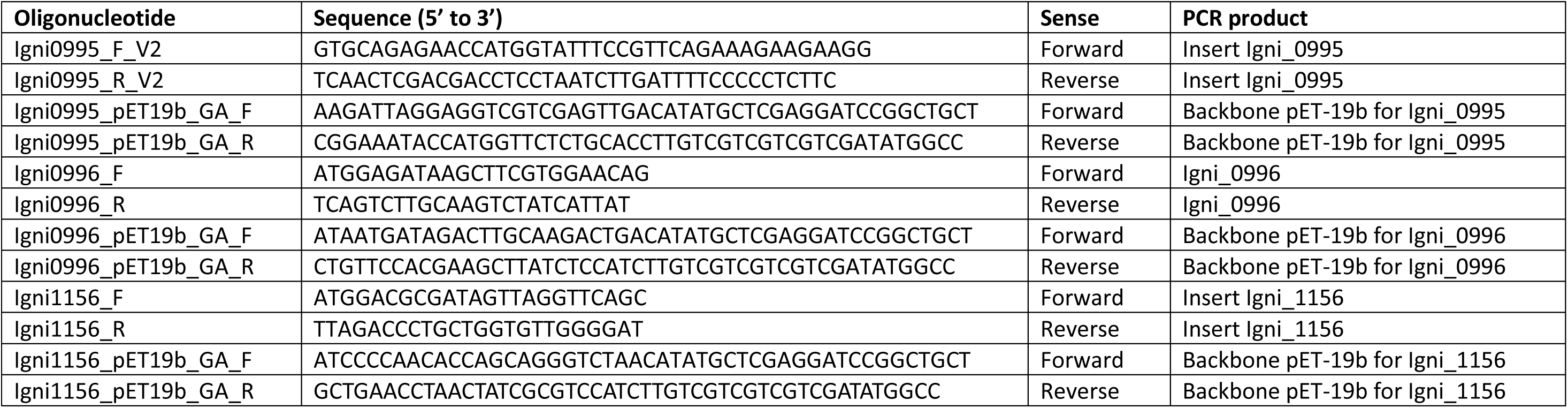
Oligonucleotides used in this study for PCR amplification of genes and plasmid backbones.

**Figure S1.**
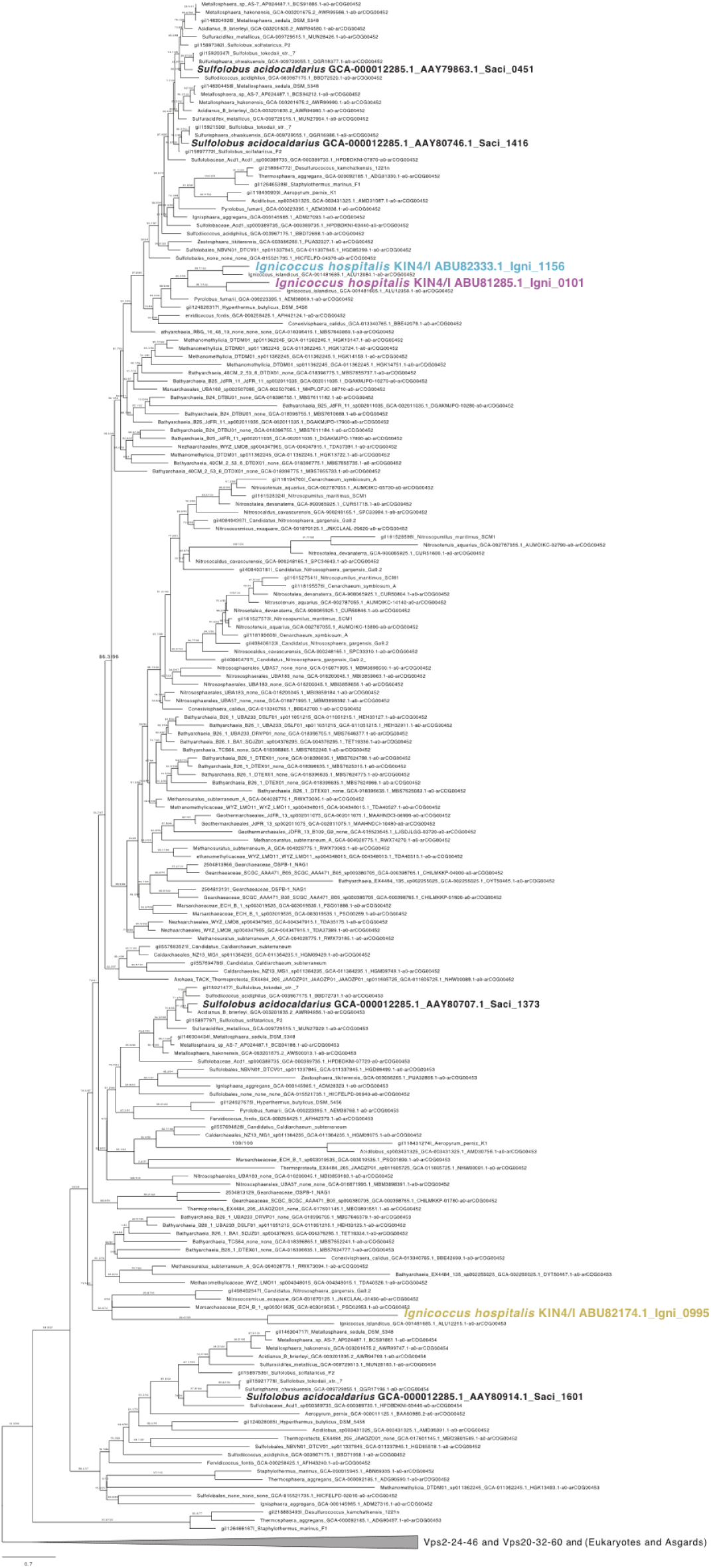
Phylogeny of CdvB, CdvB1 and CdvB2 from *Ignicoccus hospitalis*. The selected set for phylogenetic analyses contains SNF7 domain proteins of Asgard archaea and eukaryotes, e.g. homologs of Vps2-24-46 and Vps20-32-60, respectively, SNF7 domain containing proteins (Igni_0995, Igni_0101, Igni_1156) of *I. hospitalis*, as well as of homolog proteins of other archaea assigned to arCOG00454, arCOG00453 and arCOG00452. Sequences were aligned using MAFFT L-INS-I v7.407, trimmed with BMGE-1.12 and subjected to phylogenetic analyses using IQ-TREE 2 (v. 2.1.1). The tree was viewed in FigTree.

**Figure S2.**
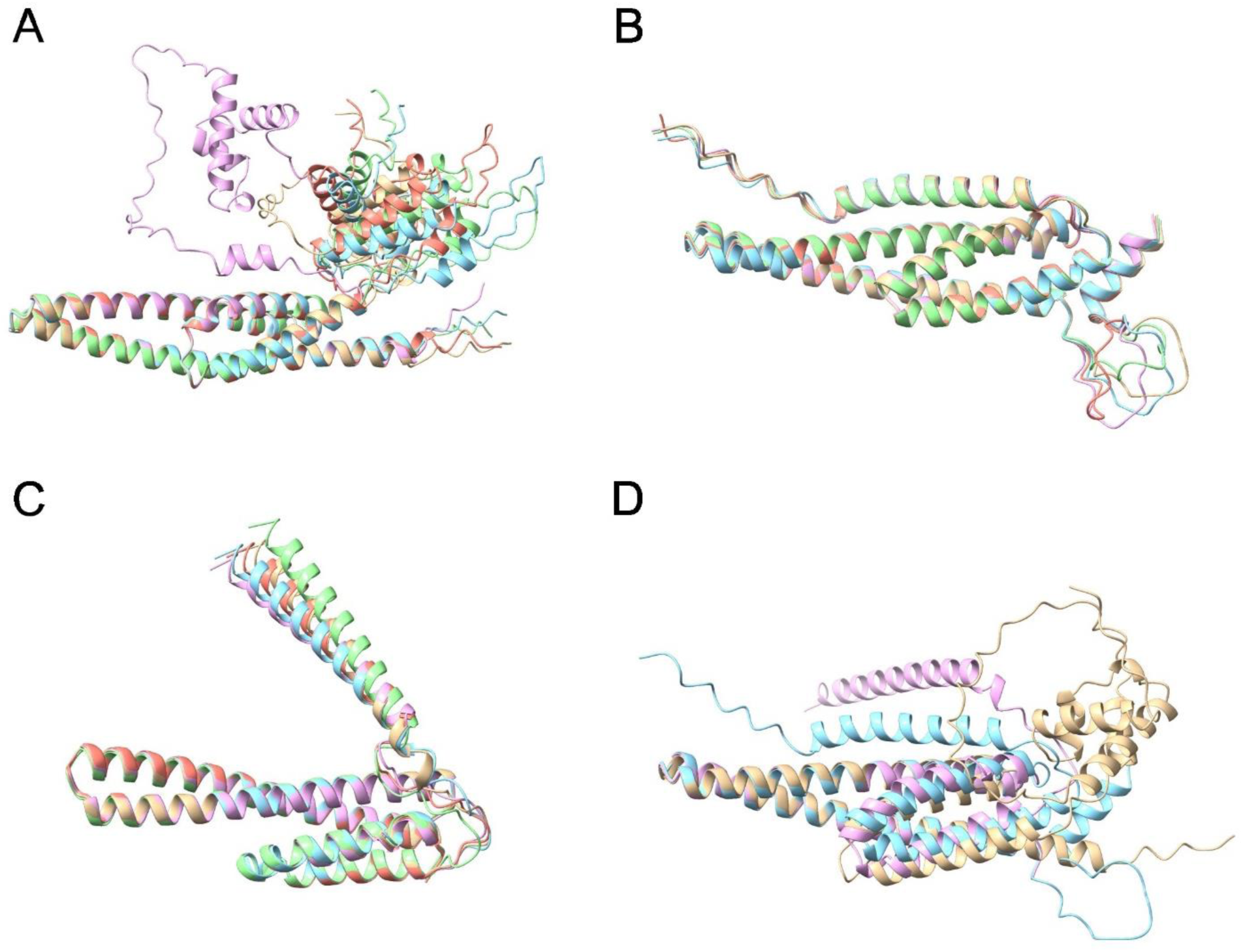
Structural prediction of *Ignicoccus hospitalis* proteins CdvB, CdvB1 and CdvB2. Structural alignment of five AlphaFold 3 models for CdvB (A), CdvB1 (B) and CdvB2 (C). For CdvB (Igni_0995) only one model (rose) differs clearly in its globular C-terminal domain. For CdvB1 (Igni_1156) and CdvB2 (Igni_0101), the models are almost identical. The most reliable model for each protein (model 0 in AlphaFold 3) was chosen for a structural alignment (D) of the protein models CdvB (gold), CdvB1 (blue) and CdvB2 (rose) with ChimeraX (v. 1.10.1). While the proteins differ structurally in their C-terminal domain, the N-terminal ESCRT-III core fold, consisting of 4 helices is highly conserved. This core fold is of crucial importance for the polymerisation of the monomers into filaments (*1*).

**Figure S3.**
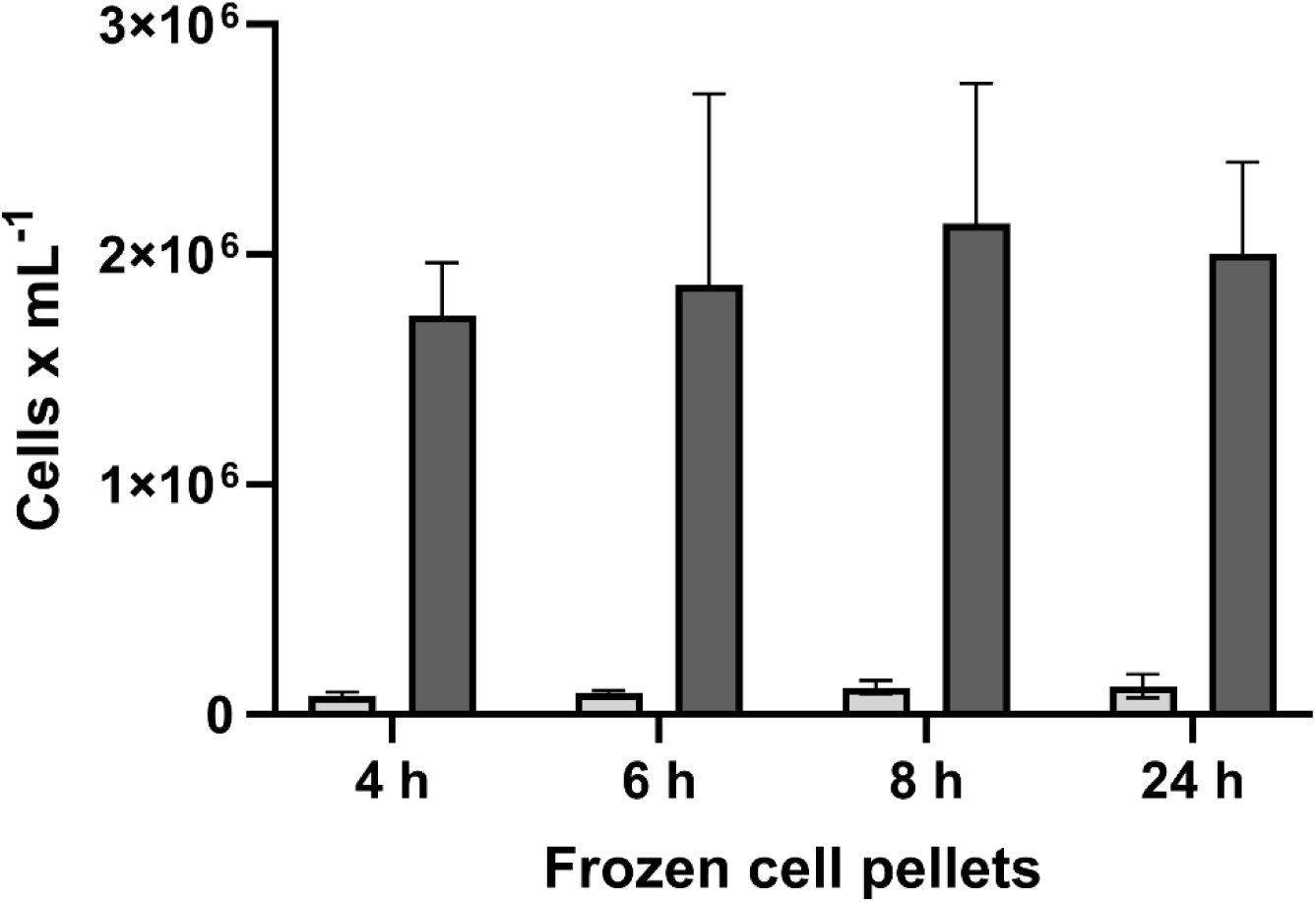
Cell viability of *Ignicoccus hospitalis* being in a state of suspended animation. Cell pellets derived from cell samples taken at different time points (4, 6, 8 and 24 h) during bioreactor cultivation were stored at −70°C and subsequently resuspended in ½ SME medium. The resuspension was used to inoculate serum bottles with 20 mL of ½ SME medium containing sulphur and H2/CO2 as gas phase. The cultures were incubated at 90°C for 24 h. The inoculation cell density (light grey bars) and final cell density after 24 h cultivation (dark grey bars) were determined by cell counting. N=3, error: SD.

**Figure S4.**
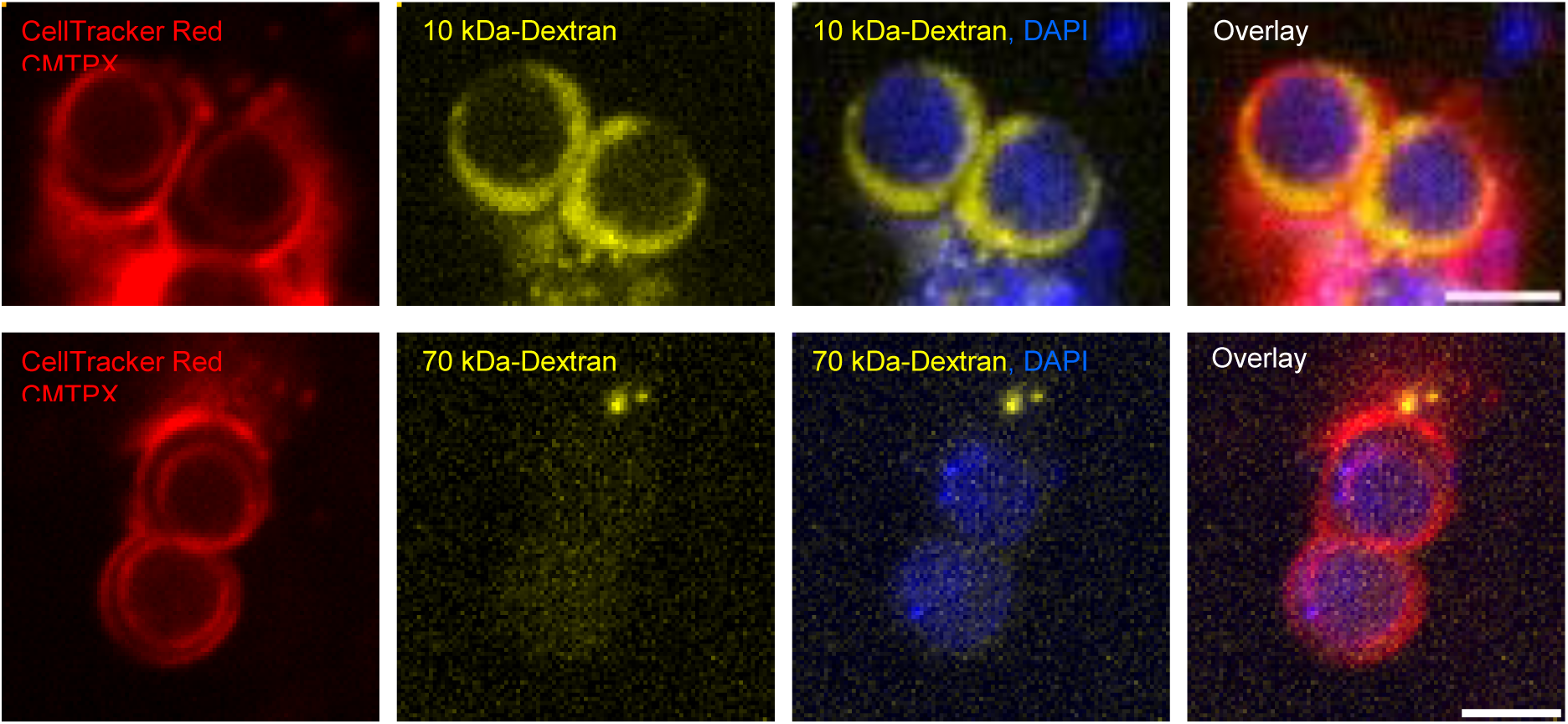
Dextran accumulation in the intermembrane compartment of *Ignicoccus hospitalis*. Membranes of *I. hospitalis* were stained with CellTracker Red CMTPX and DNA staining was performed using DAPI. Oregon Green labelled Dextran with a molecular weight of 10 kDa and 70 kDa was used, respectively. The 10 kDa-Dextran accumulates in the intermembrane compartment and the 70 kDa-Dextran is excluded from cells completely. Scale bars, 2 µm.

**Figure S5.**
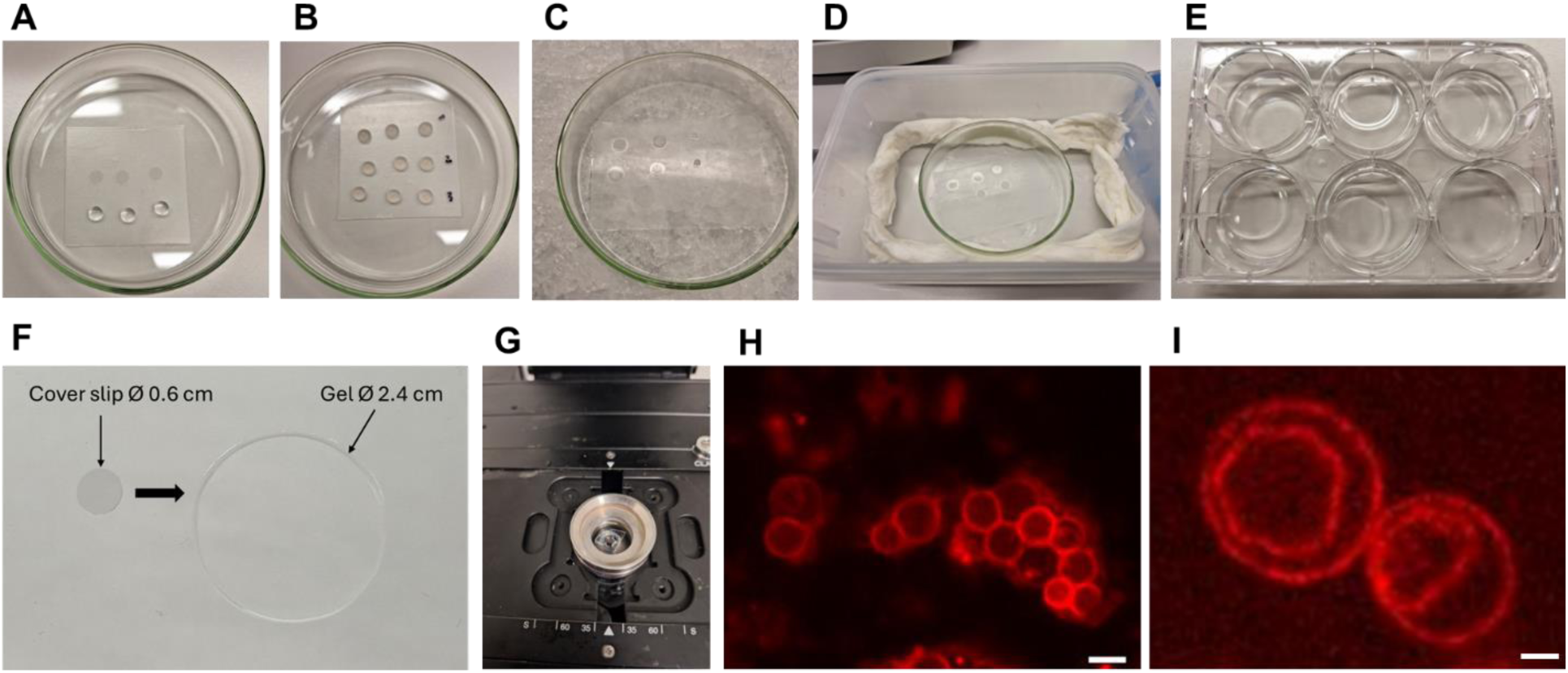
Expansion microscopy protocol developed in this study and customised for *Ignicoccus* strains. **A,** Coating of 6 mm round cover slips with poly-L-lysin solution. The cover slips were placed on parafilm in a glass petri dish. **B,** After washing the cover slips with PBS, a solution with fixed cells was dropped on the coated cover slips and incubated at room temperature. **C,** Cell solution was removed, and the cover slips were placed, slightly tilted, on 9 µL drops of gelation solution. The drops are on parafilm in a glass petri dish, which is cooled down on ice to slow down the polymerization process of the gel. **D,** Glass petri dish from C is placed in a plastic container with wet paper towels to preserve humid conditions upon closure of the box with a lid. The container was incubated at 37°C for 1 h. Afterwards the glass cover slips are placed with the gel attached into 1.5 mL reaction tubes containing denaturation buffer and incubated for 1 h at 95°C in a heating block. **E,** 6 mm gels were transferred to a 6-well plate, washed and expanded with water. **F,** 6 mm gels expanded with water to 24 mm gels, which is an expansion factor of 4. **G,** After cutting the gel in quarters, expanded cells in the gel pieces could be stained and used for immunolabeling. Imaging was performed using a spinning disk microscope and a metal ring containing a poly-L-lysin coated cover slip on the bottom. **H,** *Ignicoccus hospitalis* cells stained with Bodipy TR Ceramide, not expanded. Scale bar, 2 µm. **I,** *I. hospitalis* cells after expansion and stained with Bodipy TR Ceramide. Scale bar, 2 µm. For extended description of the protocol, see the materials and methods section.

**Figure S6.**
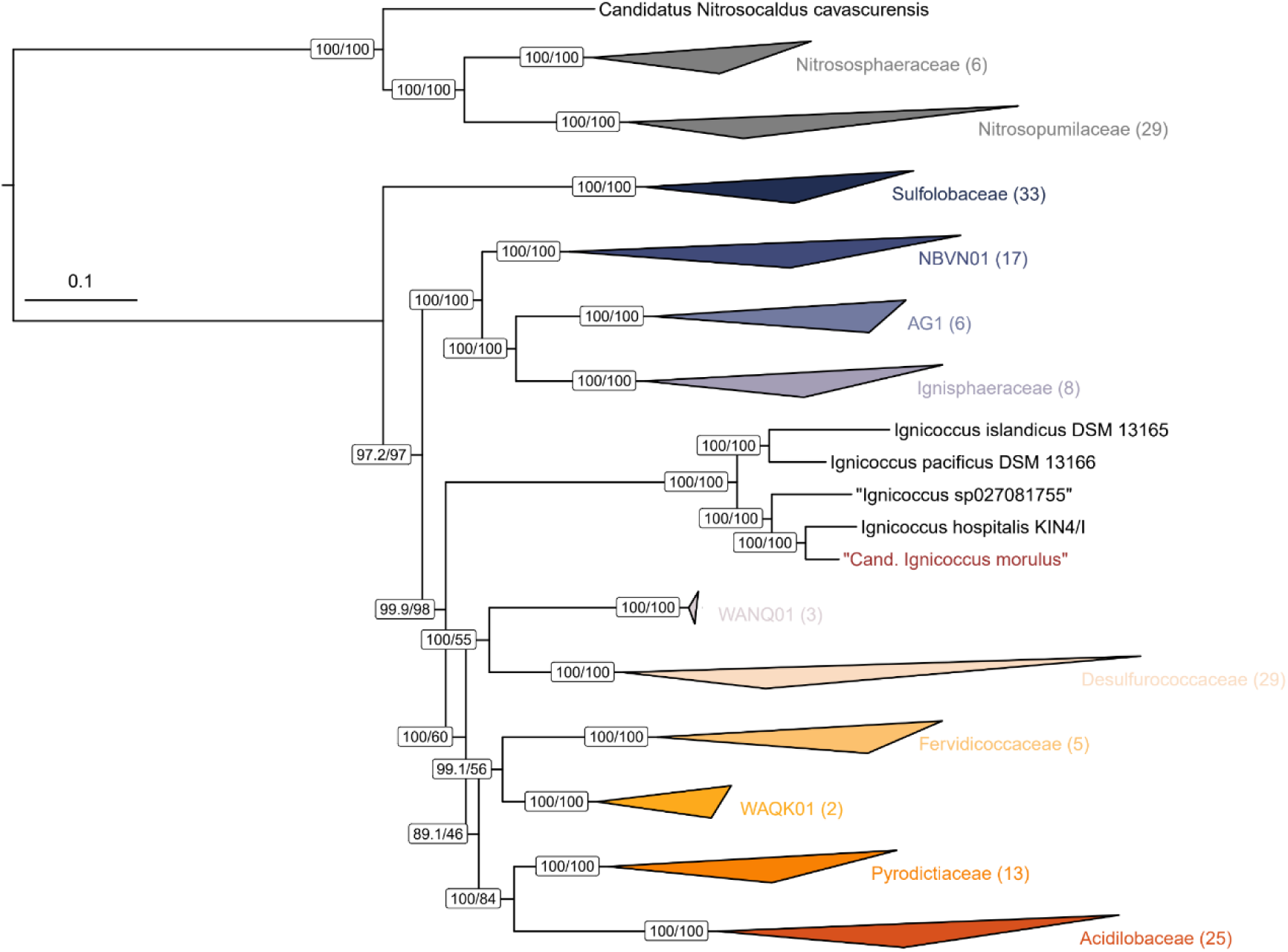
Species phylogeny of different *Ignicoccus* strains. A maximum-likelihood phylogenetic tree based on the “ar53” marker proteins (*2*) of the GTDB (*3*) and representative species of the *Sulfolobales*. Individual markers were aligned, trimmed and then concatenated. The tree was inferred in IQ-TREE (v3.0.1) with the LG+C40+F+G model with the SH-like approximate likelihood test (*4*) (left node value), and an ultrafast bootstrap approximation (*5*) (right node value), each run with 1000 replicates. The number of species represented in each clade is shown in parentheses after the taxonomic name of the clade. Scale bar: average number of substitutions per site.

**Figure S7.**
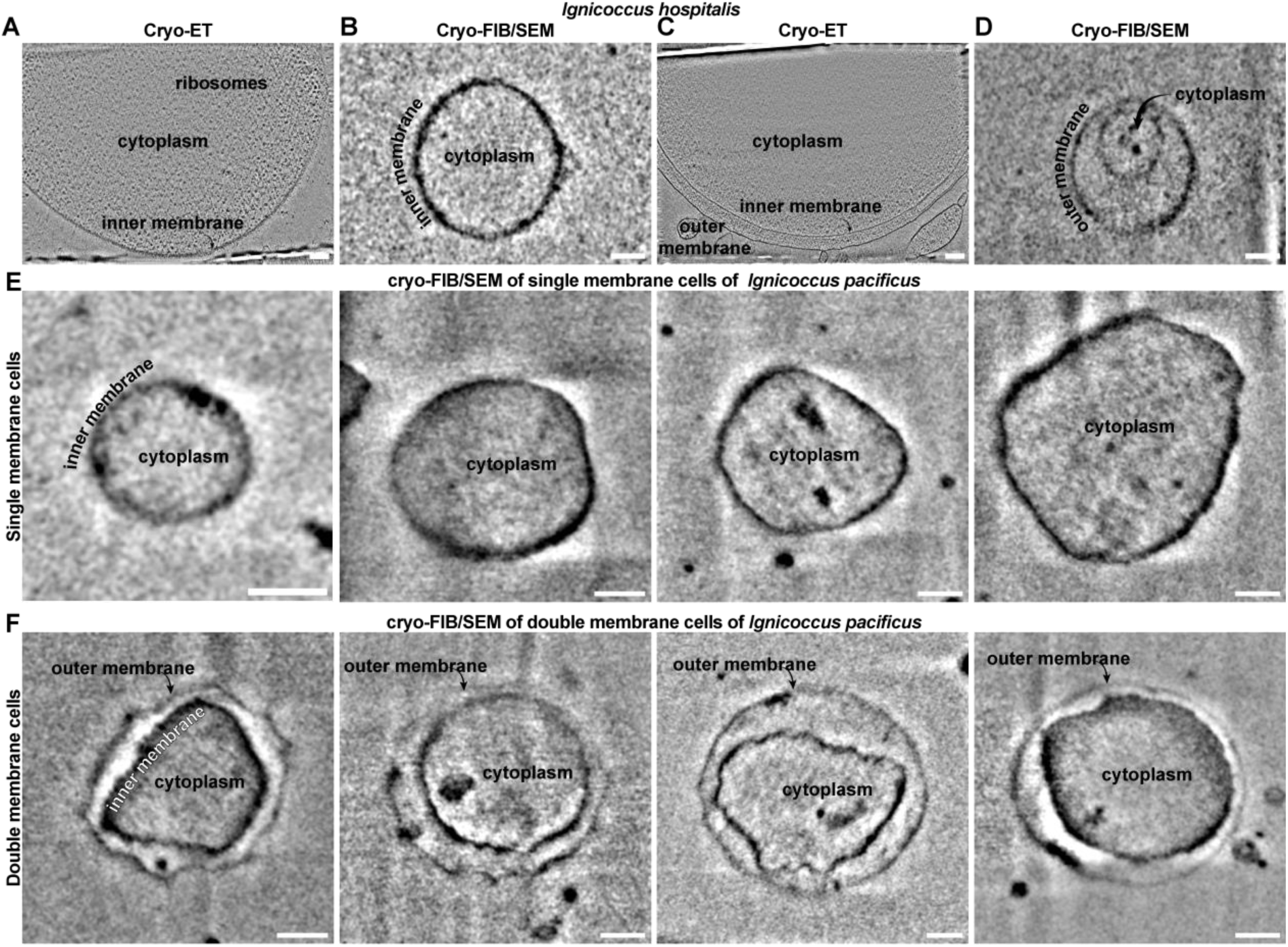
Cryogenic electron microscopy of *I. hospitalis* and *I. pacificus*. **A,** Cryo-ET slice through an *I. hospitalis* cell with a single bounding membrane. Scale bar, 100 nm. **B,** cryo-FIB-SEM image of an *I. hospitalis* cell with a single bounding membrane. Scale bar, 250 nm. **C,** Cryo-ET slice through an *I. hospitalis* cell with two bounding membranes. Scale bar, 100 nm. **D,** cryo-FIB-SEM image of an *I. hospitalis* cell with two bounding membranes. Scale bar, 250 nm. **E,** Gallery of cryo-FIB-SEM images of *I. pacificus* cells with a single bounding membrane. Scale bars, 250 nm. **F,** Gallery of cryo-FIB-SEM images of *I. pacificus* cells with two bounding membranes. Scale bars, 250 nm.

